# Multiomic network analysis reveals conserved and subtype-specific cooperative microRNA regulators in breast cancer

**DOI:** 10.1101/2025.08.07.669176

**Authors:** Eli Newby, Elijah Davis, Andrew Dhawan

**Affiliations:** Department of Cancer Sciences, Cleveland Clinic Research, Cleveland, OH, 44195, USA; Department of Neurosciences, Case Western Reserve University School of Medicine, Cleveland, OH, 44106, USA; Rose Ella Burkhardt Brain Tumor and Neuro-Oncology Center, Cleveland Clinic, Cleveland, OH, 44195, USA

**Keywords:** microRNA, Breast Cancer, Cancer Therapeutics, Gene Regulatory Networks, Protein-Protein Interaction Networks, Systems Biology

## Abstract

MicroRNAs (miRNAs) regulate gene expression through many-to-many interactions that are cooperative, context-dependent, and challenging to interpret from individual miRNA-target pairs alone. Yet, no framework simultaneously captures the cooperative, systems-level, and context-specific dimensions of miRNA regulation across the heterogeneous subtypes of a breast cancer. Here, we integrated paired miRNA and mRNA expression profiles from 1,161 breast cancers from The Cancer Genome Atlas (TCGA) with experimentally validated miRNA-target interactions and protein interaction networks to define cooperative miRNA regulatory modules in breast cancer. Community-resolved analysis of the resulting networks identified 16 functionally coherent miRNA modules associated with core cancer processes. To capture pathway-level network context beyond direct targets, we quantified the proximity of miRNA target sets to pathway proteins within a breast cancer-specific protein interaction network. This framework identified cooperative miRNA modules associated with cell cycle progression, DNA repair, PTEN/TP53-related signaling, and epithelial-mesenchymal transition. Subtype-resolved network analysis further showed that most inferred miRNA functions are strongly context dependent, not being associated in all of basal-like, HER2-enriched, luminal A, and luminal B tumors. In contrast, a limited set of pathway-level associations was conserved across subtypes. Among these, the miR-29 family (miR-29a/b/c-3p) emerged as a consistent regulator of collagen remodeling, extracellular matrix organization, PDGF signaling, and EMT. Notably, this functional conservation persisted despite subtype-specific variation in the underlying target genes. Together, these results define the cooperative miRNA modules that post-transcriptional regulate breast cancer subtypes, identify the miR-29 family as a candidate pan-subtype therapeutic target for EMT-driven metastasis, and establish a generalizable computational framework for resolving miRNA programs across heterogeneous cancers.

## 1 Introduction

MicroRNA (miRNA), short non-coding RNAs 22 nucleotides in length, are master regulators of cellular processes, ranging from development to cancer [1–4]. miRNA are the primary post-transcriptional regulators of biological systems, whose effects arise from many-to-many interactions with target transcripts. As a result of this pervasive regulation, miRNA dysregulation is consistently seen across cancer types, including breast cancer, where they perform both oncogenic and tumor suppressive roles [3, 5–7]. Because each miRNA may regulate tens to hundreds of mRNAs within the cell, and because multiple miRNA often converge on shared pathways, miRNA function is inherently cooperative and systems-level, rather than purely pairwise [8, 9]. This complexity is particularly relevant in multifactorial diseases such as cancer. For instance, the let-7 miRNA contributes to proliferation [10] and the immune response [11]. Moreover, because these interactions underlying let-7 behavior were identified separately, their combined effect has not been fully studied, and no framework is currently in place to efficiently identify this net effect of miRNA within the transcriptome. Such approaches may be leveraged to improve miRNA/siRNA-based therapeutic approaches for complex diseases such as breast cancer, which, while potentially fruitful, has currently not seen success in clinical trials [12–15].

Most studies infer miRNA function from individual miRNA-target interactions (MTIs), effectively treating each miRNA as an independent regulator [11, 16–18]. Although these approaches have identified important regulatory relationships, they are poorly suited to capture emergent properties of cooperative targeting or to differentiate between functions that are conserved across disease contexts from those that are subtype specific. In breast cancer, this limitation is particularly important because the molecular subtypes differ substantially in transcriptional state, signaling dependencies, and microenvironmental composition. Studies conducted in single cell lines or without accounting for molecular subtype generate associations that do not generalize across the heterogeneous breast cancer landscape [19, 20]. Furthermore, focusing on MTIs can only capture miRNA’s first-order effects on the transcriptome, but not higher-order effects on the proteome, resulting in a lack of systems-level information about miRNA function. No existing framework simultaneously captures the cooperative, systems-level, and context-specific dimensions of miRNA regulation — a critical gap given the clinical heterogeneity of breast cancer and the need for broadly effective therapeutic approaches.

Here, we integrate paired miRNA and mRNA expression data from 1,161 TCGA breast cancers with experimentally validated miRNA-target interactions and protein interaction networks to construct community-resolved regulatory maps of breast cancer [21, 22]. We use this framework to identify cooperative miRNA modules, infer their pathway-level functions, and compare their functional organization across the major molecular subtypes of breast cancer [23–26]. This analysis reveals both subtype-specific rewiring and a limited set of conserved pathway-level miRNA programs, exemplified by the miR-29 family, whose association with ECM remodeling and EMT is preserved across subtypes despite variation in the underlying target genes.

## 2 Results

### 2.1 Construction of a breast cancer miRNA-mRNA network identifies cooperative regulatory structure

We constructed a pan-breast cancer bipartite MTI network from 1,161 TCGA-BRCA samples (see Methods), using paired expression data and experimentally validated MTIs from miRTarBase. The final network consisted of 2,183 mRNAs, 129 miRNAs, and 5,446 validated edges (Fig. 1a,b). The stringency of experimental validation is reflected in the edge filtering: of 132,508 candidate edges based on anticorrelation alone, 127,062 (96%) were removed due to a lack of experimental evidence, ensuring the network edges reflect verified regulatory interactions rather than spurious anticorrelations. Individual miRNA connectivity ranged from 2 - 233 mRNA targets (mean 42 mRNA targets), with miR-93-5p exhibiting the broadest regulatory behavior (233 targets), consistent with its established role as a highly pleiotropic oncomiR in breast cancer [27, 28].

**Fig. 1.**
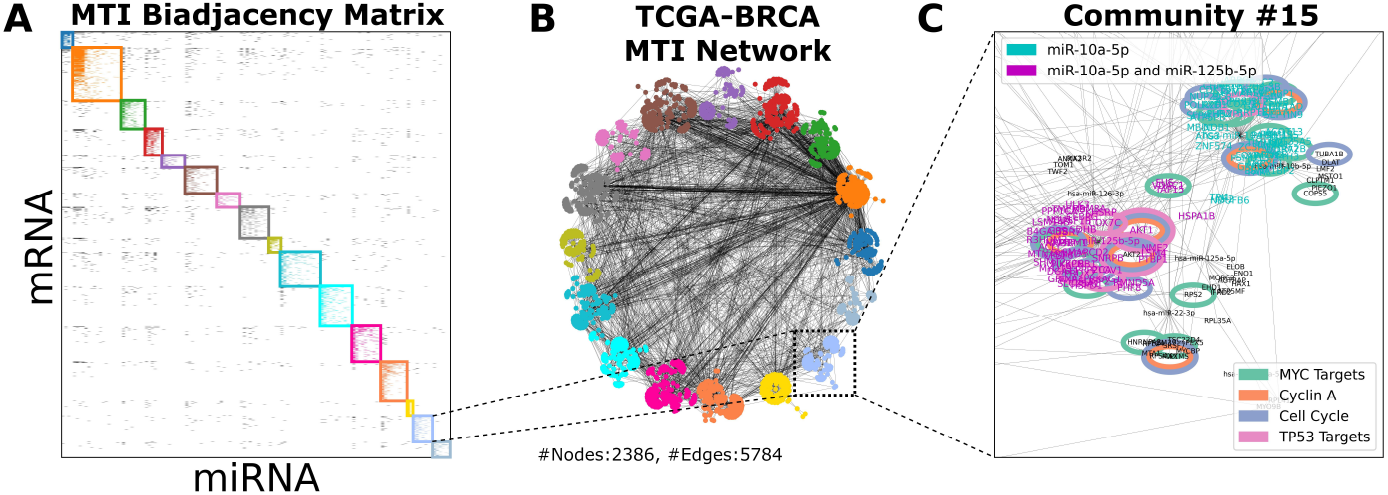
Community structure and analysis of the bipartite MTI breast cancer network. a) Biadjacency matrix of the breast cancer network organized by community. The biadjacency matrix maps the connections between the two classes of a bipartite network, in this case miRNAs and mRNAs. b) Bipartite MTI network built from breast cancer data that has been organized into individual communities. Each color represents a single community identified using the BRIM algorithm for bipartite graphs. c) The subnetwork of community 15. We indicate the genes associated with MYC Targets (*p*_*adj*_ = 4.12 *×* 10^−12^) in green, Cyclin A (*p*_*adj*_ = 9.86 *×* 10^−6^) in orange, Cell Cycle (*p*_*adj*_ = 1.22 *×* 10^−4^) in blue, and TP53 Targets (*p*_*adj*_ = 3.48 *×* 10^−4^) in pink. We illustrate the effect of driving the most influential miRNA (miR-10a-5p) with the blue labeled nodes, and the added effect of second most influential miRNA (miR-125b-5p) with the additional purple labeled nodes.

The constructed network exhibited distinct degree distributions characteristic of biological regulatory systems. mRNA nodes followed a power-law degree distribution, a scale-free topology associated with system stability, robustness, and hierarchical organization, wherein a small number of highly targeted “hub” transcripts are regulated by many miRNAs, but the majority of mRNA are regulated by a few miRNA [29–31]. miRNAs followed a log-normal degree distribution, indicating multiplicative growth of miRNA connection and resulting in most miRNA maintaining a moderate number of targets. These complementary distributions suggest that the network is organized to balance regulatory power at the miRNA level with robustness and resilience at the mRNA level, reflecting the expected biology of the system.

To characterize the cooperative structure of miRNA regulation, we analyzed the unipartite projections of the bipartite network, in which two miRNAs were connected by an edge if they shared at least one target. The average miRNA node had degree 41, indicating that a given miRNA shares a regulatory scope with, on average, 41 other miRNAs. The average clique size was 13 indicating that the breast cancer miRNA regulatory network is composed of densely interconnected cooperative groups, as opposed to isolated miRNA regulators. Conversely, the mRNA projection (in which two mRNA share an edge if co-regulated by the same miRNA) had an average degree of 179 and clique size of 32, reflecting the broad overlapping scope of miRNA-mediated post-transcriptional gene regulation across the transcriptome. These properties supported a community-based analysis of cooperative miRNA function.

### 2.2 Community structure reveals functionally coherent miRNA modules

Community detection of the bipartite network identified 16 regulatory modules (modularity *Q* = 0.56; 14 of 16 communities statistically significant at *p <* 0.05 by (q,s)-test), each representing a set of miRNAs that cooperatively target a shared gene program (Supplementary Figure 1). This modular organization, as opposed to a homogeneous or random distribution of regulatory interactions, indicates that miRNA function in breast cancer is inherently cooperative and functionally compartmentalized [32–34]. The organization of the global regulatory network into these communities is visualized in Figs. 1a and 1b, where the biadjacency matrix and network are organized by the identified communities, identified by each color. On average, we identified 210 intra-community edges in comparison to 13.8 inter-community edges emphasizing the modularity of the network. Larger communities also exhibited more extensive external connections, demonstrating a hierarchical organization where highly inter-connected modules maintain broad communication with the rest of the network. This high degree of intra- and inter-community interaction underscores the complexity and interconnectedness of the miRNA-mRNA regulatory network, where most miRNA primarily function within a densely connected cooperative module, but these modules also cooperate.

### 2.3 Functional annotation reveals coordinated miRNA co-expression driving cell cycle progression

We next sought to determine the functions of each community by examining enriched pathways and gene programs among their mRNA targets (see Methods). For instance, community 15 was significantly enriched for genes involved in cell cycle progression (Fig. 1c), where MYC Targets (*p*_*adj*_ = 4.12 *×* 10^−12^) are shown in green, Cyclin A (*p*_*adj*_ = 9.86 *×* 10^−6^) is shown in orange, Cell Cycle proteins (*p*_*adj*_ = 1.22 *×* 10^−4^) are shown in blue, and TP53 targets (*p*_*adj*_ = 3.48 *×* 10^−4^) are shown in pink. The miRNAs within this module — miR-10a/b-5p, miR-125a/b-5p, miR-126-3p/5p, miR-22-3p, and miR-320a — are thus identified as putative cooperative regulators of cell cycle progression in breast cancer. This finding is supported by prior evidence linking miR-10, miR-125, miR-126, and miR-320 to the inhibition of cell proliferation and cell cycle progression [35–38].

Notably, these miRNAs act cooperatively to target distinct transcripts within pathways associated with cell proliferation (e.g., miR-10a targets BLC6, miR-125a targets ERBB3, miR-126 targets IRS-1, and miR-320a targets AKT3). This distinct targeting, emphasizes the cooperative nature of these miRNAs in driving biological processes. This combinatorial targeting logic, wherein functionally enriched genes are regulated by multiple community members rather than a single dominant miRNA, is a defining feature of community-level regulation and demonstrates that the emergent functions of these modules cannot be inferred from any single constituent miRNA.

To further verify that enriched pathways and functions were significant beyond the first-order direct targets of the miRNA, we employed a proteomic analysis to measure higher-order effects of the miRNAs within each community [26]. Using a cancer-specific protein-protein interaction network (PPI), constructed from the STRING database combined with TCGA data (6,461 proteins and 87,759 interactions, see Methods), we assess how a miRNA’s targets affect a biological pathway’s associated proteins. We quantify the proximity of each miRNA target to the proteins of a given biological pathway using the effective graph resistance between a miRNA target and pathway protein (see Methods). Graph resistance, as opposed to the typically used graph distance, better captures the higher-order effects a dysregulated protein will have on the PPI. While most methods only compute a single shortest path between two nodes, graph resistance effectively captures the effects of both the lengths and number of paths between two nodes (e.g., two paths of length two between two points is equivalent to one path of length one). For each biological pathway, we compare the effective resistances of our miRNA targets to the effective resistances calculated on randomly chosen nodes on the PPI (with the same degree distribution) to calculate a modified Z-score indicating the size of the predicted downstream effect the miRNA’s targets may have on the target pathway proteins (Fig. 2a). That is, a more negative modified Z-score indicates that a miRNA’s targets are more closely associated with a pathway (i.e., with less resistance to the pathway targets) and will have a larger biological impact.

**Fig. 2.**
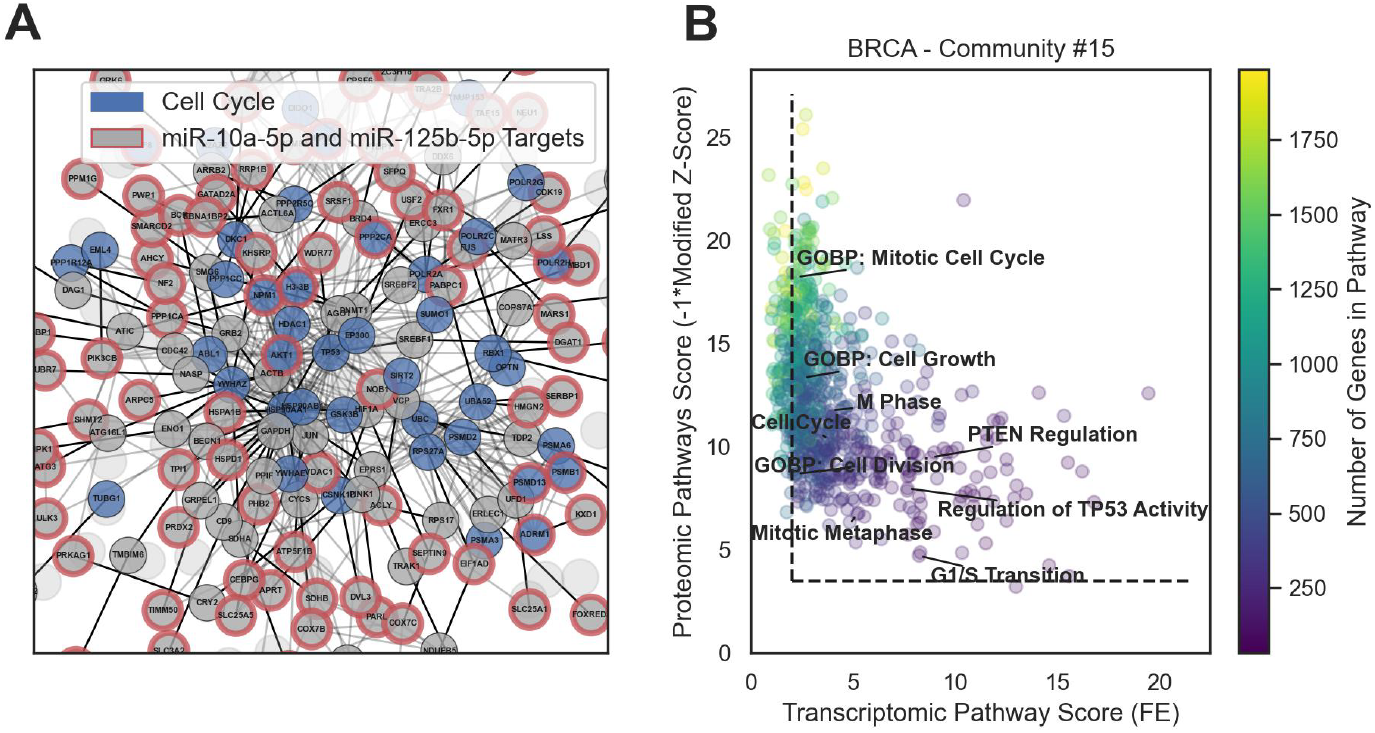
Protein-Protein Interaction (PPI) network analysis of miRNA Function. a) Sub-graph of the PPI illustrating the PPI-based analysis by showing the proximity of proteins associated with the cell cycle (blue nodes) from the proteins of genes targeted by miRNAs miR-10a-5p and miR-125b-5p (red outlined nodes) resulting in a significant modified Z-score when compared to randomly chosen PPI nodes with the same degree distribution. b) Scatter plot showing the top scoring pathways based on both the transcriptome (i.e., high fold enrichment between miRNA targets and pathways, with significance being a transcriptomic score greater than 2 (vertical line)), and the proteome (i.e., highly negative modified Z-score, indicating closer proximity to pathway proteins, with significance being a proteomic score greater than 3.5 (horizontal line)). Some significant cell cycle-related pathways are labeled on the plot. This scatter plot is also colored based on the size of the pathway to showcase the relationship between both the transcriptomic score and proteomic score with pathway size.

After calculating both a transcriptomic score (i.e., fold enrichment of miRNA targets with pathway genes) and proteomic score (i.e., modified Z-score of miRNA targets effect on pathway proteins multiplied by −1), we functionally annotate each miRNA’s predicted effect. Significant pathways are those that have both high transcriptomic and proteomic scores (Fig. 2b). Here, significance is defined as pathway fold enrichment greater than 2 and modified Z-score less than −3.5 (indicated by the dashed lines in Fig. 2b). For example, in community 15, we see that a variety of cell-cycle related pathways are significant (labeled pathways in Fig. 2b). Points in Fig. 2b are colored based on the number of genes in each pathway, thereby illuminating another quality of the transcriptomic and proteomic scores: fold enrichment is negatively correlated with pathway size while proteomic Z-score is positively correlated with pathway size. These results recapitulate that biological pathways tend to be clustered into specific subgraphs of the PPI [26, 39]. This multi-omic analysis was repeated to identify miRNA-pathway associations for all sixteen identified communities to create functional annotations of the miRNAs defining each community.

### 2.4 Global landscape of coordinated miRNA functions in breast cancer

Table 1 provides summary of the functions identified for various communities of the breast cancer network, along with the top three putative regulatory miRNA (see Supplemental Table S1 for the comprehensive table listing all identified functions for all communities). Notably, 88% of the annotated communities exhibited clear functional groupings, with 63% containing well-known breast cancer gene pathways. For each community, we list significant hallmark pathways, pathways shared across multiple communities (non-unique), and pathways unique to a single community. The associations presented in Table 1 offer a rich resource for understanding of miRNA function within breast cancer. In the following, we elaborate on three compelling associations identified using our network approach.

**Table 1.** Identified significant functions that are relevant to breast cancer and the top three miRNAs of the respective community. Identified functions are sorted into three categories: 1) Significant hallmark pathways (as defined by mSigDB), 2) Other significant pathways that are found in multiple communities, and 3) Other significant pathways that are only found in a single community. See Supplemental Table S1 for the comprehensive list of all identified functions for all communities.

| Top 3 miRNAs | Hallmark Pathways | Non-Hallmark, Non-Unique Pathways | Non-Hallmark, Unique Pathways |
| --- | --- | --- | --- |
| miR-93-5p,<br>miR-20a-5p,<br>miR-106b-5p | Apoptosis,<br>PI3K-AKT-mTOR Signaling,<br>Oxidative Phosphorylation,<br>p53 Pathway | RHO GTPase Cycle,<br>ESR Mediated Signaling | - |
| miR-15a-5p,<br>miR-195-5p,<br>miR-103a-3p | G2M Checkpoint | RHO GTPase Cycle,<br>NOTCH Signaling,<br>Cell Cycle,<br>MAPK Family,<br>WNT Signaling | - |
| miR-181(a/b)-5p,<br>miR-26a-5p | PI3K-AKT-mTOR Signaling,<br>Androgen Response | Cell Cycle,<br>MAPK Family | DNA Damage |
| miR-29(a/b/c)-3p | EMT, Myogenesis | - | PDGF Signaling |
| miR-16-5p,<br>miR-186-5p,<br>miR-148a-3p | - | WNT Signaling | Mitotic G2-G2/M Phases |
| miR-484,<br>miR-193b-3p,<br>miR-26b-5p | Mitotic Spindle | - | - |
| miR-30(a/b)-5p,<br>miR-375 | G2M Checkpoint,<br>Apoptosis | ESR Mediated Signaling,<br>WNT Signaling,<br>MAPK Family | - |
| let-7(a/b/c)-5p | DNA Repair,<br>G2M Checkpoint,<br>Oxidative Phosphorylation | ESR Mediated Signaling,<br>NOTCH Signaling,<br>WNT Signaling | S Phase |
| miR-10(a/b)-5p,<br>miR-125b-5p | Oxidative Phosphorylation | ESR Mediated Signaling,<br>WNT Signaling,<br>Cell Cycle | PTEN Regulation,<br>TP53 Regulation,<br>Mitotic G1 Phase<br>and G1/S Transition |
| miR-100-5p,<br>miR-99a-5p,<br>miR-205-5p | Oxidative Phosphorylation | DNA Replication | Hypoxia |

#### 2.4.1 The let-7-5p family of miRNA is associated with the DNA damage response

Community #13 of our network recapitulates the known roles of the let-7 family in proliferation specifically in the processing and repair of DNA. Almost all members of the let-7 family were found within the same community, implicating this miRNA family as a highly cooperative biological unit with conserved function between its members. The targets of the let-7 family were enriched for the hallmark DNA Repair and G2M Checkpoint pathways, along with pathways associated with WNT Signaling and the S Phase of the cell cycle — pathways associated with DNA replication and DNA damage response. Our results highlight the let-7 family as a negative regulator of these pathways by targeting multiple subunits of the 20S proteosome (PSMA2, PSMB2) as well as CUL1, a G1/S transition associated gene. The let-7 family’s control over the cell cycle is then specialized towards cell cycle checkpoints and DNA damage repair by the targeting of the RAD21, SMC1A, and RFC2 transcripts.

#### 2.4.2 miR-10-5p and miR-125-5p cooperatively regulate PTEN and TP53, two essential pathways in breast cancer

Along with miRs-10(a/b)-5p and miR-125-5p being drivers of the cell cycle, our network analysis also revealed a novel cooperative association between these miRNAs and regulation of pathways associated with two key genes in breast cancer: PTEN and TP53. Within community #15, PTEN pathway activity was found to be negatively regulated by miR-10a/b-5p targeting GATAD2A, PSMB1, PSMD11, and PSMD13 and miR-125-5p targeting the ADRM1 and AKT1 transcripts. TP53 activity is regulated by miR-10a/b-5p targeting CDK2, and GATAD2A. Likewise, miR-125-5p targets AKT1, PIP4K2C, PPP2CA, and PRKAG1 within the TP53 pathway. The gene TAF15, a negative regulator of p53, is negatively regulated by both miR-10-5p and miR-125-5p. These associations are displayed in figure 3. Although miR-10a/b-5p and miR-125-5p are predicted to target mostly different genes within these pathways, the combined effect of these genes enhances the regulation of these two pathways. Furthermore, gaining an understanding of how these miRNAs interact is important for the prediction of higher-order effects such as the existence of competing endogenous miRNAs. Understanding these first-order cooperations and their higher-order effects highlight the strength of this community-based analysis.

**Fig. 3.**
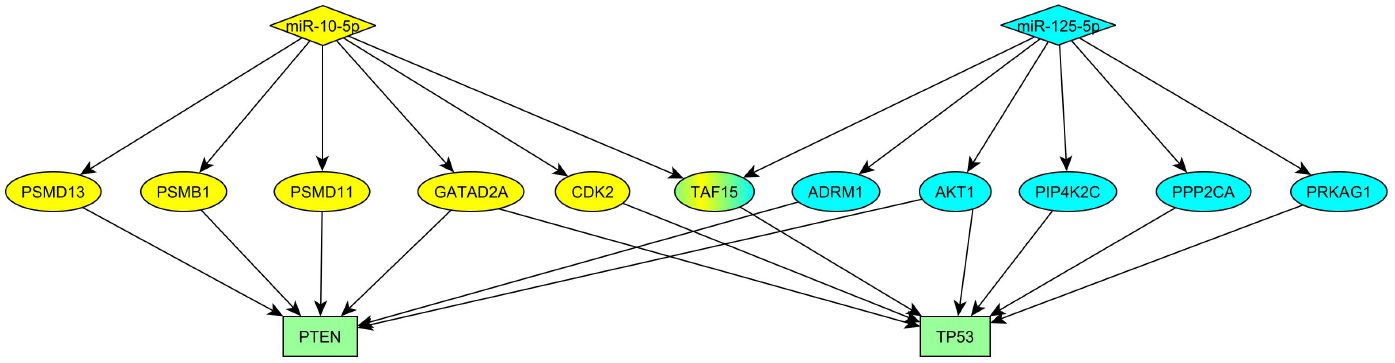
Visual representation of the miRNAs in community #15, miR-125-5p and miR-10-5p, and their post-transcriptional regulation of genes related to the PTEN and TP53 pathways.

#### 2.4.3 The miR-29-3p family was identified to regulate pathways responsible for breast cancer metastasis, and may be a therapeutic target

In community #9, the miR-29 family was predicted to be a strong negative regulator of processes related to collagen, the extracellular matrix (ECM), and the epithelial-to-mesenchymal transition (EMT). This finding consolidates previous independent observations linking the miR-29 family to TGF-*β* expression to ECM and collagen production[40], and further linking TGF-*β* production to EMT [41, 42]. Furthermore, this miRNA family was also identified to be uniquely associated with PDGF signaling pathways, which is heavily linked to metastasis of breast cancers. Our analysis demonstrates that it is the combination of the three miRNAs miR-29a/b/c-3p that results in these functions. These three miRNAs share only seven mRNA targets and thus each target different components of these pathways, highlighting a naturally occurring combinatorial targeting approach to pathway regulation, which could be exploited for therapeutic benefit.

Further exploring combinatorial targeting, we define and calculate a miRNA combination effectiveness score (see Methods) to assess the potential downstream effects of therapeutic cocktails of miRNA, and which act synergistically. Table 2 presents the top three miRNA cocktails (i.e., the miRNA pairs expected to most strongly downregulate a given pathway) for five cancer-related functions. Among these combinations, we show that the combination of endogenous miR-29b-3p and miR-29c-3p are expected to be highly cooperative in downregulating the EMT pathway, earning an effectiveness score much higher than any other miRNA combination. Thus, combinatorial targeting of the miR-29 family — in particular the miR-29b/c-3p pair, which achieved the highest effectiveness score across all tested pathways, represents a compelling therapeutic strategy for disrupting EMT-driven metastasis in breast cancer.

**Table 2.** Most effective pairs of miRNA for driving select pathways relevant to breast cancer. Effectiveness is calculated as the absolute value of the sum of each miRNA’s target percentages multiplied by the combined miRNA target set’s PPI-based Z-score.

| Pathway | Best miRNA Combinations | Effectiveness |
| --- | --- | --- |
| DNA Repair | let-7a-5p, let-7b-5p | 1.86 |
|  | let-7b-5p, let-7c-5p | 1.79 |
|  | let-7b-5p, let-7f-5p | 1.72 |
| EMT | miR-28b-3p, miR-29c-3p | 3.55 |
|  | miR-29a-3p, miR-29b-3p | 2.78 |
|  | miR-182-5p, miR-29b-3p | 2.55 |
| ESR Mediated Signaling | miR-185-5p, miR-30a-5p | 2.13 |
|  | miR-93-5p, let-7a-5p | 2.07 |
|  | miR-106b-5p, miR-93-5p | 2.01 |
| MAPK Family | miR-101-3p, miR-200c-3p | 1.27 |
|  | miR-15b-5p, miR-101-3p | 1.24 |
|  | miR-26a-5p, miR-30a-5p | 1.24 |
| PI3K-AKT-mTOR Signaling | miR-106b-5p, miR-20a-5p | 1.75 |
|  | miR-17-5p, miR-20a-5p | 1.26 |
|  | miR-20a-5p, miR-181a-5p | 1.23 |

### 2.5 Subtype-specific miRNA-mRNA networks reveal context dependent regulation of gene expression

A central challenge in breast cancer biology is identifying molecular regulators that are therapeutically actionable across the heterogeneous molecular subtypes. Our subtype-specific network analysis directly addresses this challenge by revealing which miRNA regulatory programs are conserved versus divergent across the Basal-like, HER2-Enriched, Luminal A, and Luminal B subtypes. We constructed bipartite MTI networks and context-specific PPIs for each breast cancer subtype using the same methodology as the original, global breast cancer network (see Methods). Supplementary Table S2 summarizes the properties of each of these four constructed networks. All subtype networks maintained similar structural characteristics as the global network, including comparable ratios of mRNAs to miRNAs (approximately 19:1) and edges to nodes (approximately 2.4:1), as well as consistent power-law mRNA and log-normal miRNA degree distributions. This structural consistency supports the validity of our community-based approach when applied to each breast cancer subtype.

#### 2.5.1 Subtype-specific analysis reveals conserved and divergent miRNA functions

Across breast cancer molecular subtypes, the overlap in miRNA targets and associated pathways were analyzed. While certain miRNAs exhibited high overlap in mRNA targets across subtypes (Fig. 4a), a more striking finding was the overall low functional overlap amongst most miRNAs, with only 24 unique functions identified in all four subtypes (Fig. 4b). Here we focus on annotations that are unique between a miRNA and function to restrict our analysis to miRNA functions that are specific to a miRNA community and not functions that may be shared amongst multiple communities. Approximately 30% of the identified functions were only found in a single subtype of breast cancer. Of the 24 overlapping unique functions, only the miR-29 family (miR-29a/b/c-3p) showed consistently high functional conservation across subtypes (Fig. 4c). To further analyze all potentially conserved miRNA functions, table 3 lists the (non-unique) hallmark pathway-miRNA pairs that are conserved across all four molecular subtype networks for pathways relevant to breast cancer (see Supplemental Table S3 for the complete list of conserved hallmark pathway-miRNA pairs for all hallmark pathways). For a comprehensive summary of the functional associations for every community in all four molecular subtype networks see Supplemental Tables S4-S7. The discrepancy between target and functional similarity underscores that most miRNA do not have specific, conserved function across contexts and that functional annotation derived from our network approach may provide a more accurate representation of context-dependent miRNA importance than mere target overlap.

**Table 3.**
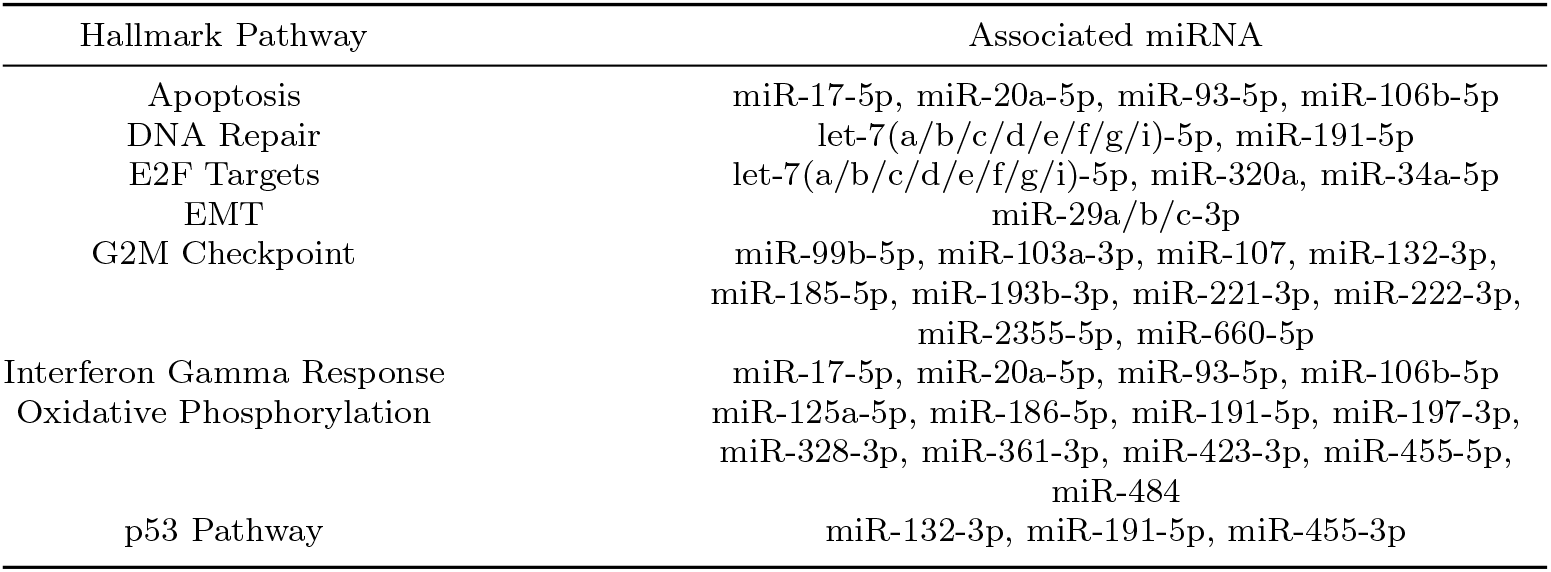
A subset of hallmark pathways that are relevant to breast cancer and the miRNA that are consistently associated with these pathways across all four breast cancer molecular subtype networks. See Supplemental Table 3 for the consistent associations between all hallmark pathways and miRNA.

| Hallmark Pathway | Associated miRNA |
| --- | --- |
| Apoptosis | miR-17-5p, miR-20a-5p, miR-93-5p, miR-106b-5p |
| DNA Repair | let-7(a/b/c/d/e/f/g/i)-5p, miR-191-5p |
| E2F Targets | let-7(a/b/c/d/e/f/g/i)-5p, miR-320a, miR-34a-5p |
| EMT | miR-29a/b/c-3p |
| G2M Checkpoint | miR-99b-5p, miR-103a-3p, miR-107, miR-132-3p,<br>miR-185-5p, miR-193b-3p, miR-221-3p, miR-222-3p,<br>miR-2355-5p, miR-660-5p |
| Interferon Gamma Response | miR-17-5p, miR-20a-5p, miR-93-5p, miR-106b-5p |
| Oxidative Phosphorylation | miR-125a-5p, miR-186-5p, miR-191-5p, miR-197-3p,<br>miR-328-3p, miR-361-3p, miR-423-3p, miR-455-5p,<br>miR-484 |
| p53 Pathway | miR-132-3p, miR-191-5p, miR-455-3p |

**Fig. 4.**
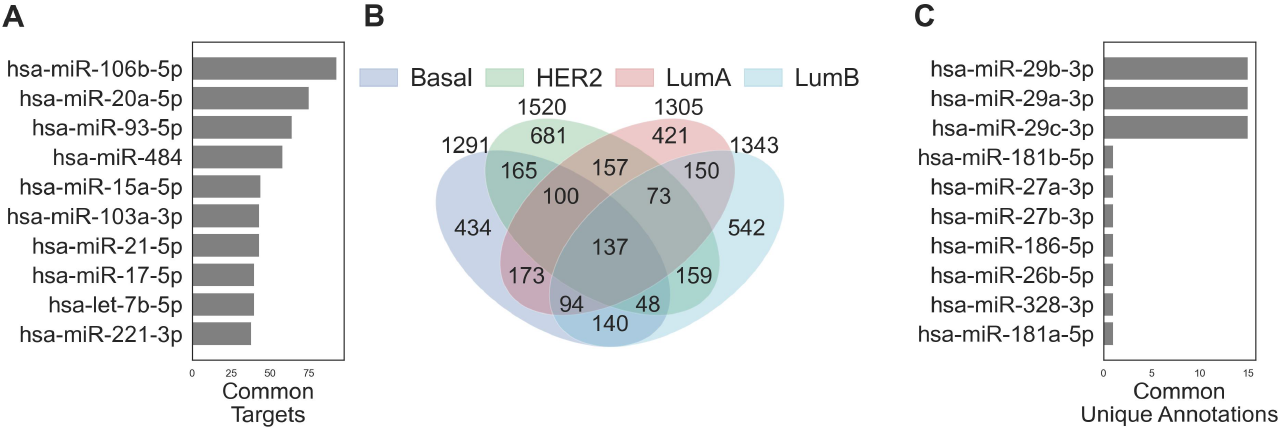
Comparisons of miRNA across the four breast cancer subtype networks. a) Bar chart of the miRNAs with the ten largest sets of common targets. b) Venn diagram showing the overlap of the unique functions for each of the communities in the subtype networks. b) Bar chart of the miRNAs with the ten largest sets of common miRNA-function associations.

#### 2.5.2 The miR-30 family displays context-dependent function amongst breast cancer molecular subtypes

While miR-30 emerged as a key regulator of the multiple important cancer pathways within the global breast cancer network, this miRNA family displays distinct functions across breast cancer subtypes. Among the non-unique pathway associations, the miR-30 family is a driver of many housekeeping pathways, but because these pathways are non-unique, they are not the only miRNAs associated with these pathways, so the system may be more robust to the effects of introducing a therapeutic targeting miR-30. The miR-30 family did not have any unique functions common across all four subtypes and only one function common to three subtypes (Xenobiotic Metabolism was associated with miR-30 in the Basal and both Luminal subtypes). Most of miR-30’s unique functions were identified in a single subtype, with the identified functions not being biologically similar. In the Basal subtype, miR-30 was found to be associated with calcium ion binding. In the HER2 subtype, miR-30 was found to be associated with exonuclease activity. In the Luminal A subtype, miR-30 was found to be associated with KRAS upregulated genes and the RHO GTPase cycle. In the Luminal B subtype, miR-30 was found to be associated with signal transmission and DNA synthesis. These disparate functions demonstrate that miR-30 has many potential functions, but these functions are highly context-dependent.

#### 2.5.3 miR-181 is an oncogenic regulator of PI3K-AKT-mTOR Signaling in Basal-like breast cancers

The miR-181 family also highlights the need for context-dependent analysis of miRNA function, as a global approach may make incorrect generalizations. In the global breast cancer network miR-181 was found to be a driver of the PI3K-AKT-mTOR Signaling pathway and DNA damage pathways, recapitulating miR-181’s identified role as an oncomiR. Analysis of the breast cancer subtype networks reveals that the miR-181 family does not have any consistent associations across subtypes. This analysis also reveals that the associations between miR-181 and PI3K-AKT-mTOR Signaling and DNA damage are only present in the Basal-like subtype of breast cancer. The miR-181 family has fewer and more marginal annotations in the other subtypes, emphasizing that the results from the global network were likely a consequence of only the Basal cells in the sample. Thus, miR-181 if used as a cancer therapeutic, would only be expected to be effective for Basal-like breast cancers and to a lesser extent in other subtypes.

#### 2.5.4 The miR-29 family is a pan-subtype regulator of EMT and extracellular matrix remodeling

The miR-29 family (miR-29a/b/c-3p) emerged as the most functionally conserved miRNA family across all four breast cancer subtypes, consistently occupying the highest-degree position within the collagen/ECM/EMT community in each subtype network (Fig. 2d). This conservation is particularly striking because the precise mRNA targets of the miR-29 family show substantial subtype-specific variation (maximum Jaccard similarity of miRNA targets between any pairs of subtypes is 0.36 between Luminal A and Luminal B subtypes), demonstrating that functional conservation can be maintained through distinct molecular mechanisms in different cellular contexts. The miR-29 family therefore represents a functionally conserved, rather than merely mechanistically conserved, pan-subtype regulator of EMT, a key driver of metastatic breast cancer. This finding has direct therapeutic implications: a miRNA therapeutic targeting the miR-29 family would be predicted to inhibit EMT-driven processes across all four molecular subtypes, regardless of the specific mRNA targets mediating this effect in each subtype.

## 3 Discussion

Here, we move beyond pairwise miRNA-target analyses, and establish that a cooperative module — distinct from individual miRNA — is a more accurate functional unit of post-transcriptional gene regulation in breast cancer, and we establish that functional conservation can be maintained across heterogeneous breast cancer subtypes despite changes in target genes. Three findings stand out for their biological and translational significance. First, the breast cancer miRNA-mRNA network is organized into functionally coherent communities that encode cooperative gene programs, including cell cycle regulation and EMT, not predictable from individual miRNA analyses. Second, the miR-29 family is the only miRNA family in our analysis whose functional association with EMT and PDGF is preserved across all four molecular subtypes of breast cancer, despite variation in its subtype-specific target genes. Third, our subtype-resolved networks reveal that miRNA functional identity is shaped more by cellular context than by target gene overlap, with miR-30 and miR-181 serving as striking examples of a miRNA whose functional associations vary markedly across subtypes despite consistent network prominence.

While individual links between miR-29, ECM, and EMT have been previously reported [43–45], our network approach consolidates these observations, and, crucially, reveals that this relationship is conserved across heterogeneous breast cancer molecular subtypes. The miR-29 family possesses a robust, fundamental role in these processes, suggesting it is a strong candidate pan-subtype therapeutic target for inhibiting EMT-driven metastasis. Our analysis also revealed that additional established EMT regulators (e.g., p53 and MYC pathways) were not consistently targeted by the same miRNA across breast cancer subtypes, further highlighting the uniquely broad therapeutic advantage of targeting the miR-29 family. Further, our analysis has shown that miR-10a-5p and miR-125b-5p, two miRNAs that have previously been linked to a number of functions in breast cancer via target-based analyses [46–48], are both key putative regulators of cell cycle genes and regulators of TP53 and PTEN activity, helping to refine wide-ranging prior experimental evidence into synergistic functions for these miRNA.

Individual networks constructed for each of the four breast cancer molecular subtypes allowed us to directly interrogate the context-dependence of miRNA function, addressing a major challenge in developing therapeutics for this heterogeneous disease. While some miRNAs have conserved target gene sets, unique functional conservation is less common, reinforcing the concept that a miRNA’s role is profoundly shaped by its cellular milieu. The behaviors of the miR-29 family and miR-30 act as instructive examples: both are identified as key regulators in each of the four breast cancer molecular subtypes, but the functional associations of the miR-29 family are consistent across subtypes whereas the functional associations for miR-30 varied significantly. This analysis also distinguishes the miR-29 family from the miR-17 family, which showed high conservation in both targets and functions, but the miR-17 family’s conserved functions suggest a potentially narrower therapeutic utility and may be more representative of a housekeeping miRNA family.

While effective for identifying regulatory, context-specific functions of miRNA, our analysis does present some limitations. An important direction for future work will be to extend the community detection framework to allow for overlapping community membership. This would better capture the known pleiotropy of biological pathways and the multi-community regulatory roles of individual miRNAs, which we see in interconnectedness of our identified communities. Strict thresholding and multiple layers of validation (e.g., (q,s)-test for communities, Fisher’s exact test for pathways, thresholding on modified Z-Scores for the proteome) applied throughout the analysis of these networks increases confidence but may further restrict the results. A secondary limitation of this approach is its reliance on available biological data, limiting discovery to pre-existing pathways, without considering the biological context in which those pathways were discovered. This reliance on experimental data may also result in unidentified off-target effects for less well-studied miRNA. Lastly, our approach relies on the use of bulk RNA-sequencing data to construct the miRNA-mRNA interaction networks, limiting the usage of this approach to largely homogeneous populations of cells, such as tumor cores. Thus, while this study provides robust and powerful computational predictions of global miRNA effects and functions, we acknowledge that experimental validation of our putative miRNA targets is crucial.

Collectively, this work establishes a network community framework for dissecting the cooperative, context-dependent regulatory logic of miRNA in cancer. By moving beyond pairwise miRNA-target analyses to model the emergent properties of miRNA modules, we reveal biologically coherent regulatory programs. By focusing on the biological context, we identify the unique pan-subtype association of the miR-29 family with EMT and extracellular matrix remodeling. This approach is thus able to identify novel relationships and filter previous global results that are invisible to conventional approaches. Furthermore, the framework is generalizable: the same pipeline can be applied to any cancer type or complex disease for which paired miRNA-mRNA expression data exist, offering a scalable strategy for mapping cooperative miRNA function across biological contexts. More broadly, these findings underscore that miRNA therapeutic targeting may be most effective when designed at the level of cooperative modules rather than individual miRNAs, a principle with direct implications for the rational design of miRNA-based therapeutics in breast cancer and beyond.

## 4 Methods

### 4.1 Data Availability

The code used to complete this analysis is available at https://doi.org/10.5281/zenodo.17109250[49]. Gene and miRNA expression data was accessed (Feb 2025) and downloaded through the NIH Genomic Data Commons (GDC) TCGA Pan-Cancer Atlas, as reprocessed by Hoadley et al[22]. This dataset comprises *ex vivo* tumor samples from 1,161 patients who provided informed consent as part of The Cancer Genome Atlas Project, with all procedures reviewed and approved by local Institutional Review Boards. Breast cancer molecular subtypes were classified using the TCGAquery subtypes function in the TCGABiolinks R package [50, 51]. For BRCA, this function classifies cases into one of four subtypes Basal-like, Her2, Luminal A, or Luminal B — based on the molecular features of these subtypes [21]. Experimentally verified human miRNA-target interactions were obtained from miRTarBase, downloaded in October 2024 [52, 53]. Protein-protein interaction networks were constructed from STRING-db data downloaded March 2026 [54].

### 4.2 Network construction

The following four-step process is used to construct our networks:

1. Pare down the TCGA-BRCA data to the samples that have both mRNA expression values and miRNA expression values. We notate mRNA expression values of a single patient sample *k* as 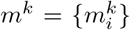, where *i* indexes the mRNAs within the sample. Similarly, we notate miRNA expression values for sample 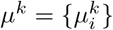.
2. Identify nodes:
  a. a. Find the set of mRNAs *M* that are in the intersection of the mRNAs in the top 50% of expression values for each sample. This can be represented as

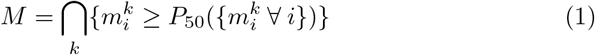

where *P*_*n*_ is a function that returns the nth percentile of the input set.
  b. Find the set of miRNAs *µ* that are in the intersection of the miRNAs in the top 50% of expression values for each sample. The calculation of *µ* is equivalent to equation 1, except 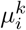 is used instead of 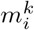.

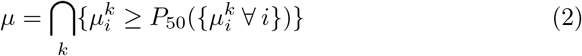
  c. The nodes of the Network 𝒩 are the union of the two sets 𝒩 = *M* ⋃ *µ*.
3. Identify edges:
  a. Identify all possible miRNA-mRNA pairs from the identified nodes, which will become the set of potential edges: *E* = *M × µ*
  b. Calculate the spearman correlation *r*_*s*_(*m*_*i*_, *µ*_*i*_) between mRNA *m*_*i*_ and miRNA *µ*_*i*_ expression values for every potential edge, and discard all positive correlations

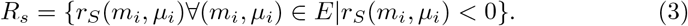

Note: We only look at pairs with negative correlations because we are interested in the miRNAs that cause mRNA degradation.
  c. The edges *ε* of the network are the correlations that are in the bottom 50% of the negative correlations (i.e., the most negative correlations).

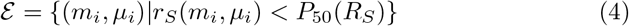
4. Filter the identified edges through miRTarBase. Within the set of experimentally-verified miRNA-target interactions in miRTarBase, we keep all identified edges that have experimental backing (e.g., HITS-CLIP, Western Blot, Microarray, ChIP-Seq).

Filter threshold percentiles from 50 (inclusive) to 100 (exclusive) in increments of 5 were tested, and these alternate threshold values did not change network structure (Supplemental Figure S2).

### 4.3 Network community detection

To identify communities with coordinated miRNA function on mRNA targets within the constructed network, we applied the Bipartite Recursively Induced Modules (BRIM) algorithm, implemented in Python https://github.com/genisott/pycondor [55–57]. Briefly, the BRIM algorithm is a spectral optimization method that recursively optimizes the modularity of the bipartite network, resulting in the identification of distinct communities. The BRIM algorithm exploits the structure of the bimodularity matrix **B**. *B*_*ij*_ = *A*_*ij*_ − *p*_*ij*_, where **A** is the biadjacency matrix of the network, and *p*_*ij*_ represents the probability of an edge between nodes *i* and *j* under a defined null model. The algorithm optimizes the modularity function:

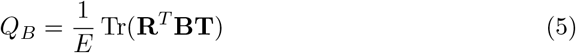

where *E* is the number of edges in the network, **R** is the initial community membership matrix for the mRNAs, and **T** is the initial community membership matrix for the miRNAs. Tr denotes the trace of the matrix. BRIM recursively calculates, from the initial configuration, the set of miRNAs and mRNAs that maximize *Q*_*B*_. BRIM does not guarantee finding the community structure that is the global maximum *Q*_*B*_, but does guarantee that a local maximum for *Q*_*B*_ will be identified from our initial community configuration. In our case, the initial community configuration was defined using the function `greedy_modularity_communities` in the networkx library, which uses a greedy algorithm to identify communities maximizing modularity in a unipartite network.

### 4.4 Network community validation

Network community statistical significance was validated using the (q, s)-test implemented in Python https://github.com/skojaku/qstest[58], which compares each community’s modularity to communities of similar size in randomly generated networks with the same degree distribution. For our analysis, we generate 1000 random networks with the same structure. A community was deemed statistically significant if it achieved a p-value less than 0.05 when comparing the modularity of the network community to the random communities.

### 4.5 Fisher’s exact test and fold enrichment to identify community function

To identify potential functions (e.g., pathways, gene ontologies) of the genes within each community we employed Fisher’s exact test and fold enrichment. Any fold enrichment greater than 1 was taken to indicate overrepresentation compared to the background. P-values were adjusted for multiple testing. Statistical significance for identified functions was defined as an adjusted *p <* 0.05. To identify the set of functions for a community, we determine which significant functions (i.e., functions with *FE >* 1 and *p*_*adj*_ *<* 0.05) are unique to this community.

We curated a custom set of gene pathways by integrating those from mSigDB [59] and the DAVID Bioinformatics functional annotation tool [60]. Our analysis focused on pathways and ontologies that were indicative of functional roles for miRNA in breast cancer. All human genome pathways form mSigDB were used. From the DAVID Bioinformatics functional annotation tool, we utilized the following gene sets: `UP_KW_BIOLOGICAL_PROCESS`, `UP_KW_MOLECULAR_FUNCTION`, `GOTERM_BP_DIRECT`, `GOTERM_MO_DIRECT`, `BBID Pathways`, `BIOCARTA Pathways`, `KEGG_PATHWAY`, `REACTOME_PATHWAY`, and WIKIPATHWAYS. Although the two subsets had some overlap of terms, using distinct gene lists enhanced robustness of our analysis.

### 4.6 Protein-protein interaction network construction

The context-specific PPI’s were constructed using data from v12.0 of STRING database. Human protein interactions were downloaded in March 2026, and interactions with a combined score greater than 0.7 were included in the final network. The included proteins were then made to be context-specific by filtering the PPI to only contain proteins associated with the mRNA that were found to be in the top 50% of all TCGA samples (*M*).

### 4.7 Graph effective resistance-based analysis of miRNA targets in the PPI

The effective graph resistance measures the resistance between two nodes if every edge is assumed to be one ohm resistor. Here, to find the proximity of each miRNA’s targets to a given biological pathway, we perform the following steps:

1. For each miRNA target, we find the smallest effective resistance between a miRNA target and every pathway protein, and then average these resistances. 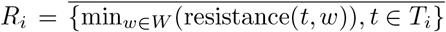, where *T*_*i*_ are the targets of miRNA *µ*_*i*_, and *w* are the protein in pathway *W*. The bar represents the average of the set.
2. Repeat step 1 for 100 random miRNA target sets and pathway protein sets that have the same degree distributions.
3. Calculate the modified Z-score of the average resistance value using the median *med* and median absolute deviation *MAD* calculated from the values in step 2, where the modified Z-score is defined as: *Z* = 0.6745(*R*_*i*_ − *med*)*/MAD*

This modified Z-score is then used to assess the proximity of a set of miRNA’s targets to a biological pathway with a modified Z-score less than −3.5 indicating significance. We use a modified Z-score in our analysis because the graph effective resistance is prone to outliers.

### 4.8 microRNA combination effectiveness score

The miRNA combination’s effectiveness score is defined as follows:

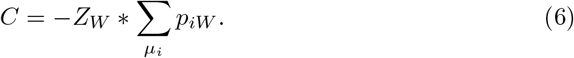

Where, *Z*_*W*_ represents the modified Z-score of comparing the set of miRNAs’ targets to the biological pathway *W*, *µ*_*i*_ represents miRNA *i*, and *p*_*iW*_ represents the percentage of the biological pathway that is in miRNA *i*’s targets.

## Author Contributions

EN: Conceptualization, Methodology, Software, Investigation, Formal Analysis, Validation, Visualization, Writing — Original Draft. ED: Investigation, Writing — Review and Editing. AD: Conceptualization, Resources, Data Curation, Writing — Review and Editing, Supervision, Funding Acquisition.

## Acknowledgments

The results shown here are in whole or part based upon data generated by the TCGA Research Network: https://www.cancer.gov/tcga. A.D. acknowledges the American Academy of Neurology Clinical Research Training Scholarship. This work was supported by the National Cancer Institute, National Institutes of Health [5T32CA094186-23 to E.N.].

