## Supplemental Material for "Multiomic network analysis reveals conserved and subtype-specific cooperative microRNA regulators in breast cancer"

Figures

Fig. S1.

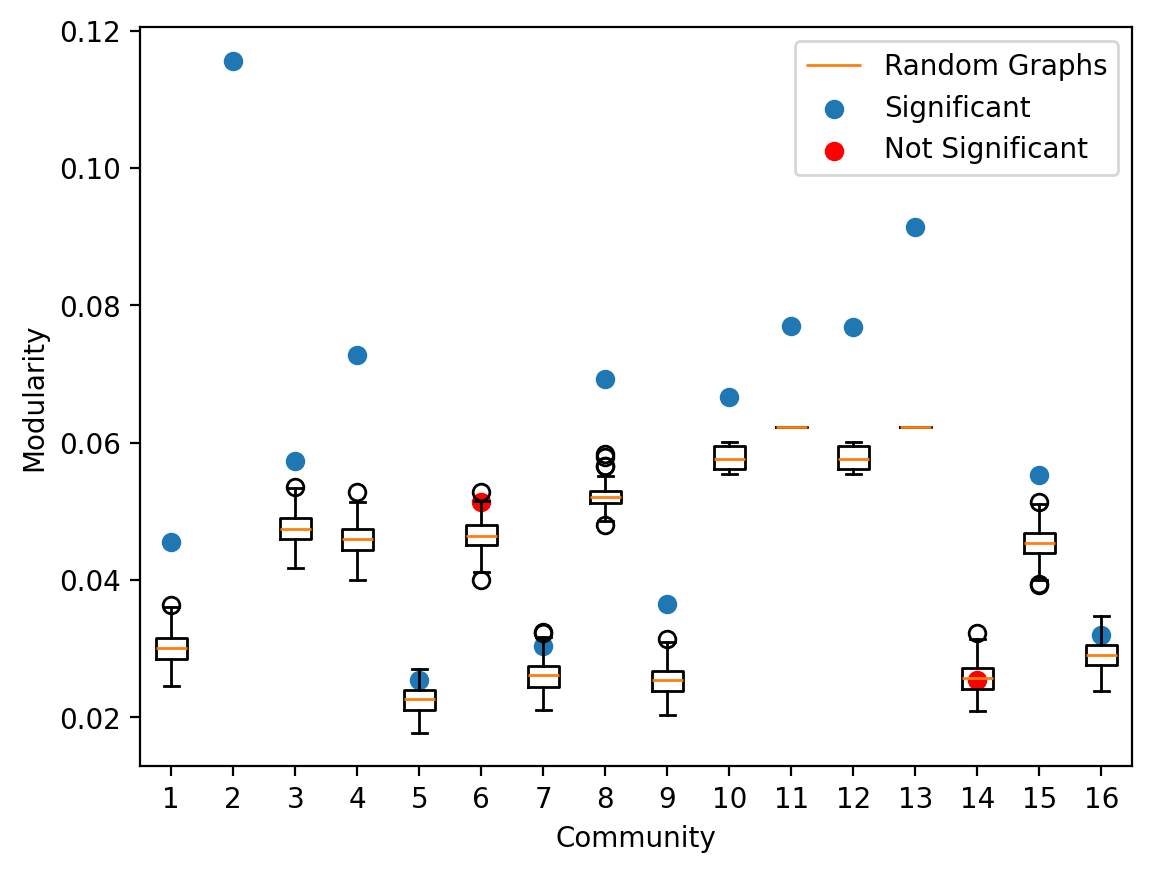

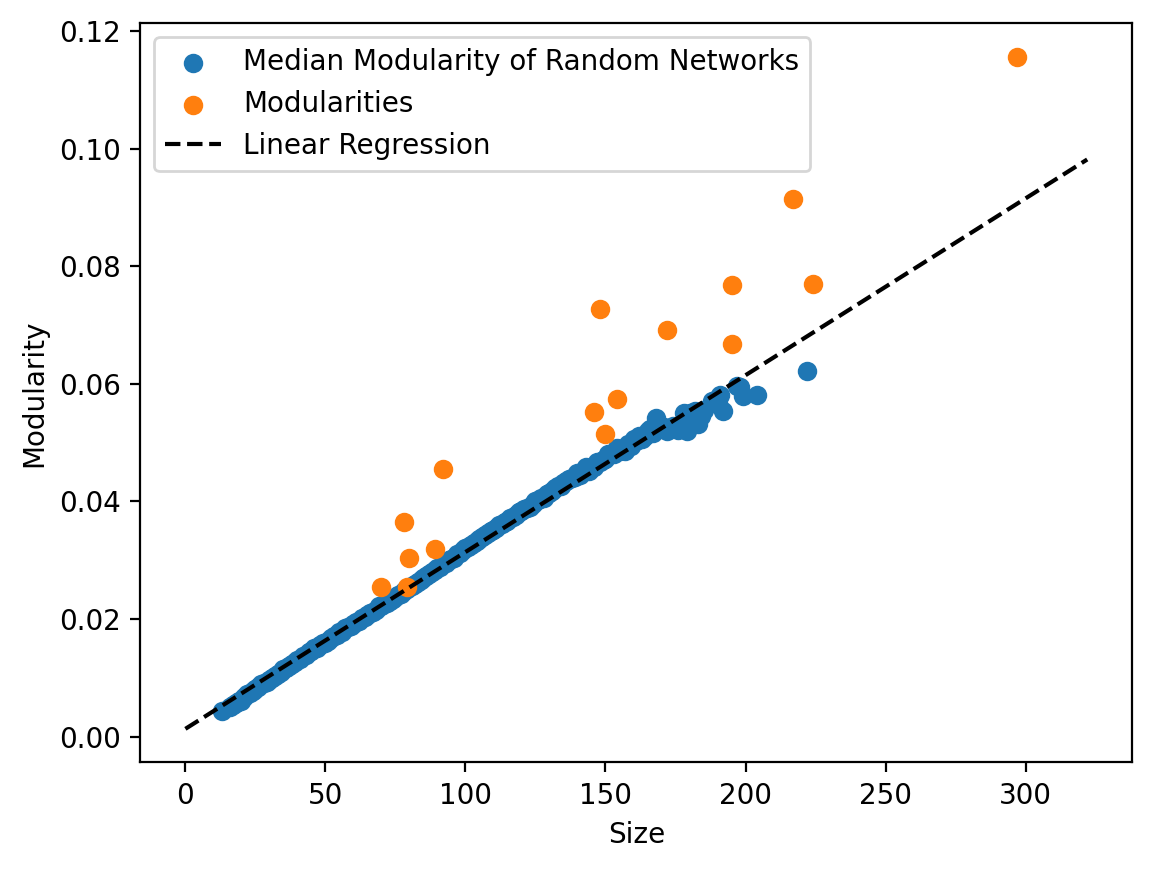

a) To assess the validity of a community, we use the (q,s)-test with 1000 randomly generated networks. We compare the modularity of communities in the BRCA network (circular points) to communities of similar size (i.e., +/- 10 nodes) in the random networks (box plots). Communities 2 and 15 do not have a boxplot because there were no communities of similar size in the randomly generated networks. In all but two cases, the community is statistically significant (indicated by a blue point) in terms of having a larger modularity than random communities. b) Scatter plot of the size of communities vs the modularity of those communities, which is used to assess the communities that don’t have any similarly sized communities in the random networks (i.e., communities 2 and 15). The orange points indicate the modularities of the BRCA communities, and the blue points represent the modularities of the communities identified in the randomly generated networks, which showcase a linear relationship. To extrapolate the size-modularity relationship for larger sizes, we calculate the linear regression of the random data (dashed line), which shows that communities 2 and 15 have modularities higher than randomly expected.

Fig. S2.

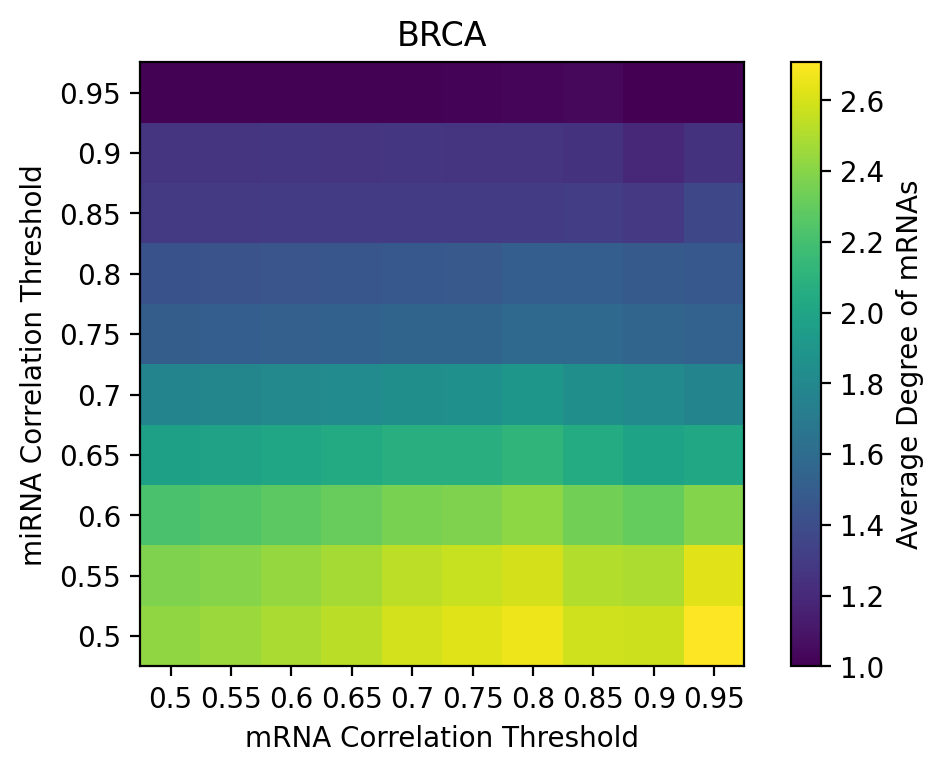

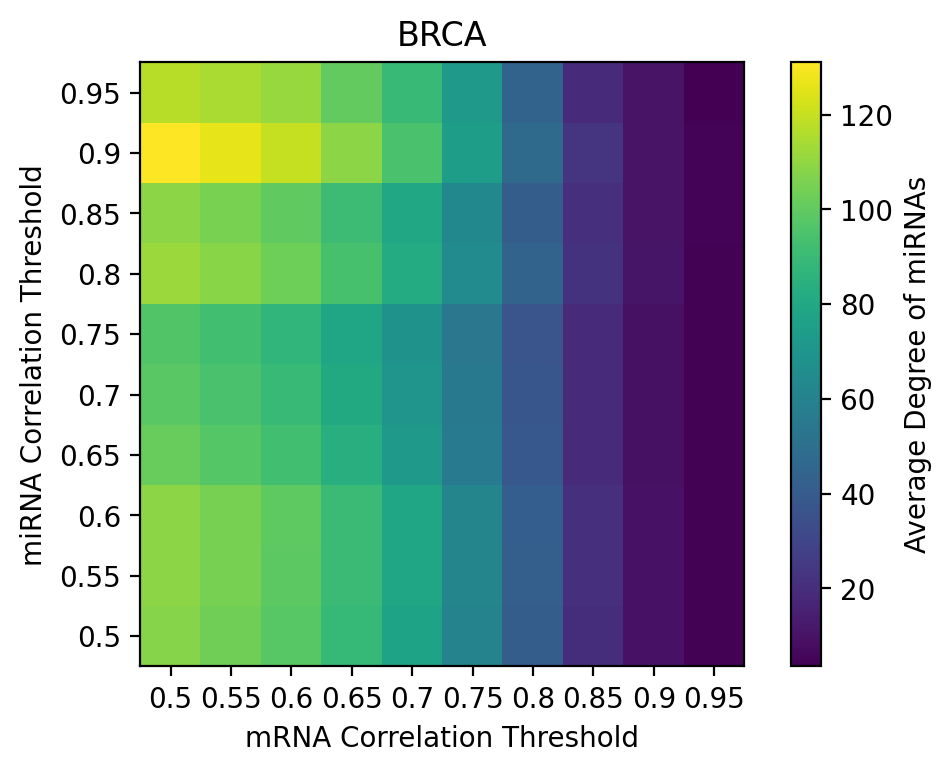

The average degree of the mRNAs (left) and the miRNAs (right) at different tested percentile thresholds. While there is some variation, in general, the average degree of the mRNAs is mostly dependent on the miRNA threshold, and the average degree of the miRNAs is mostly dependent on the mRNA threshold. This indicates that the number of identified connections mostly depends on the number of identified mRNAs or miRNAs, but that increasing the number of mRNAs does change the number of connections for other mRNAs in the network, and similarly for the miRNAs. Thus, the threshold itself does not change the structure of the network, and only affects the number of nodes that are in the network.

Tables

**Table S1.** Identified functions and top three miRNAs for each community in the global breast cancer network with relevant citations.

| **Top 3 miRNAs** | **Hallmark Pathways** | **Non-Hallmark, Non-Unique Pathways** | **Non-Hallmark Unique Pathways** |
| --- | --- | --- | --- |
| miR-27(a/b)-3p, miR-128 3p | - | Histone Modifying Activity, Transcription Factor Binding, Chromatin Binding, Post Transcriptional Protein Modification | Cis-regulatory Region Sequence-specific DNA Binding |
| miR-93-5p, miR-20a-5p, miR-106b-5p | Protein Secretion(1–3), Adipogenesis, Apoptosis(4), PI3K-AKT-mTOR(4), Oxidative Phosphorylation, Fatty Acid Metabolism, p53 Pathway(5, 6) | RHO GTPase Cycle, Membrane Trafficking, Immune Protein Signaling(7–9), ESR Mediated Signaling(6, 10, 11), Transferase, Transcription | Transcription Repressor Complex, Ubiquitin Binding |
| miR-92a-3p, miR-455-3p, miR-25-3p | - | Vesicle Mediated Transport(12, 13), Transcription, Cell Adhesion, Cell-Cell Junction, RTK Signaling | Calcium Ion Binding |
| miR-15a-5p, miR-195-5p, miR-103a-3p | MYC Targets V1, G2M Checkpoint | RHO GTPases, NOTCH Signaling, Cell Cycle(14), Cell Cycle Checkpoints, M Phase, Vesicle Mediated Transport, Deubiquitination, Spindle, MAPK(14, 15) Family, WNT Signaling(14, 15), Cell-Cell Junction, Immune Protein System | HIV Life Cycle(16–18) |
| miR-140-3p, miR-524-3p, miR-146a-5p | - | Cell Adhesion(19), Immune Protein Signaling, Cell Cycle | - |
| miR-155-5p, miR-221-3p, miR-24-3p | Not Significant | Not Significant | Not Significant |
| miR-320a, miR-34a-5p, miR-365a-3p | - | Transcription, Cell Adhesion, Chromatin Binding, Nervous System Development | Sequence Specific DNA Binding |
| miR-181(a/b)-5p, miR-26a-5p | PI3K-AKT-mTOR Signaling(20), mTORC1 Signaling, MYC Targets V1, Androgen Response | Cell Cycle, Immune Protein Signaling(21), Transcription, MAPK Family(22), RTK Signaling | DNA Damage, Bacterial Infection Pathways, Cytoplasmic Stress Granule, NTRKs, Toll-like Receptor Cascades |
| miR-29(a/b/c)-3p | EMT(23, 24), Myogenesis(25, 26) | RTK Signaling, Microtubule Cytoskeleton, Cell Cycle | Collagen/Extracellular Matrix(24, 27, 28), MET, Scavenger Receptors, PDGF Signaling |
| miR-16-5p, miR-186-5p, miR-148a-3p | Heme Metabolism | Nonsense Mediated Decay, Translation, Splicing, Cell Cycle, WNT Signaling(29, 30) | Cellular Senescence(31, 32), Mitotic G2-G2/M Phases |
| miR-484, miR-193b-3p, miR-26b-5p | MYC Targets V1, Mitotic Spindle | Translation(33, 34), Splicing, Anchoring Junction, Vesicle Mediated Transport | Small Ribosome(33, 35), rRNA, Mitotic Spindle, Sm-like Protein Family |
| miR-30(a/b)-5p, miR-375 | MYC Targets V1, Unfolded Protein Response, G2M Checkpoint, Xenobiotic Metabolism, UV Response Up, E2F Targets, Apoptosis, Complement, mTORC1 Signaling, Heme Metabolism, Glycolysis | ESR Mediated Signaling(36), WNT Signaling(37), Immune Protein Signaling, Cell Cycle, ALK Signaling, MAPK Family(38), RTK Signaling, Autophagy | Double Stranded RNA Binding, B-Cell Receptor, CLRs |
| let-7(a/b/c)-5p | MYC Targets V2, MYC Targets V1, Unfolded Protein Response, Oxidative Phosphorylation, E2F Targets, DNA Repair(39–41), UV Response Up, mTORC1 Signaling, Adipogenesis, G2M Checkpoint | Translation, Nonsense Mediated Decay, NOTCH Signaling, Cell Cycle(42–44), ESR Mediated Signaling, WNT Signaling, Immune Protein Signaling | ATP Dependent Activity Acting on RNA, Acetyltransferase Complex, Single Stranded DNA Binding, Nucleotide Excision Repair, S Phase, MITF-M |
| miR-21-5p, miR-21-3p | Not Significant | Not Significant | Not Significant |
| miR-10(a/b)-5p, miR-125b-5p | MYC Targets V1, Oxidative Phosphorylation, Adipogenesis, mTORC1 Signaling, E2F Targets | Nonsense Mediated Decay, Splicing(45), Translation, ESR Mediated Signaling, WNT Signaling, Cell Cycle | PTEN Regulation(46), TP53 Regulation(47), Mitotic G1 Phase and G1/S Transition |
| miR-100-5p, miR-99a-5p, miR-205-5p | MYC Targets V1, Oxidative Phosphorylation | DNA Replication, Cell Cycle(48) | Hypoxia(49, 50), Electron Transport |

Table S2. Properties of the datasets and networks constructed for each of the breast cancer subtypes.

| **Subtype** | **Samples** | **Nodes** | **mRNAs** | **miRNAs** | **Edges** |
| --- | --- | --- | --- | --- | --- |
| Basal | 192 | 3067 | 2914 | 153 | 7069 |
| HER2 | 82 | 3778 | 3592 | 186 | 9429 |
| LumA | 562 | 3050 | 2898 | 152 | 7257 |
| LumB | 209 | 3136 | 2973 | 163 | 7674 |

**Table S3**, Hallmark pathways and the miRNA that are associated with this pathway across all four breast cancer subtype networks. For each subtype, we identify which miRNAs target which of the hallmark pathways (i.e., the genes for the hallmark pathway are enriched in the same community as a given miRNA), and then find which of these miRNA-hallmark pathway pairs is found in all four of the subtype networks.

| Hallmark Pathway | miRNAs |
| --- | --- |
| Adipogenesis | let-7a/b/c/d/e/f/g/i-5p, miR-17-5p, miR-20a-5p, miR-93-5p, miR-106b-5p |
| Apical Junction | miR-660-5p |
| Apoptosis | miR-17-5p, miR-20a-5p, miR-93-5p, miR-106b-5p |
| DNA Repair | let-7a/b/c/d/e/f/g/i-5p, miR-191-5p |
| E2F Targets | let-7a/b/c/d/e/f/g/i-5p |
| Epithelial Mesenchymal Transition | miR-29a/b/c-3p |
| G2M Checkpoint | miR-103a-3p, miR-107, miR-132-3p, miR-660-5p |
| Heme Metabolism | miR-103a-3p, miR-107, miR-660-5p |
| Hypoxia | miR-101-3p, miR-423-3p, miR-484 |
| Interferon Gamma Response | miR-17-5p, miR-20a-5p, miR-93-5p, miR-106b-5p |
| Mitotic Spindle | miR-26b-5p, miR-27a/b-3p, miR-103a-3p, miR-107, miR-128-3p, miR-132-3p, miR-660-5p |
| mTORC1 Signaling | miR-26b-5p, miR-27a/b-3p, miR-30a/b/c/d/e-5p, miR-101-3p, miR-125a/b-5p, miR-126-3p, miR-128-3p, miR-132-3p, miR-140-5p, miR-181a/b-5p, miR-186-5p, miR-191-5p, miR-328-3p, miR-374a-3p, miR-423-3p, miR-484. miR-660-5p |
| MYC Targets v1 | let-7a/b/c/d/e/f/g/i-5p, miR-29a/b/c-3p, miR-30a/b/c/d/e-5p, miR-125a/b-5p, miR-126-3p, miR-132-3p, miR-140-5p, miR-145-5p, miR-181a/b-5p, miR-186-5p, miR-191-5p, miR-328-3p, miR-338-3p, miR-423-3p, miR-484 |
| Myogenesis | miR-29a/b/c-3p |
| Oxidative Phosphorylation | miR-125a-5p, miR-186-5p, miR-191-5p, miR-197-3p, miR-328-3p, miR-423-3p, miR-455-3p, miR-484 |
| p53 Pathway | miR-132-3p, miR-191-5p, miR-455-3p |
| Protein Secretion | miR-17-5p, miR-20a-5p, miR-93-5p, miR-106b-5p |
| UV Response Down | miR-132-3p |
| UV Response Up | miR-30a/b/c/d/e-5p |

**Table S4.** Identified functions and top three miRNAs for the communities in the Basal subtype network.

| **Top 3 miRNAs** | **Hallmark Pathways** | **Non-Hallmark, Non-Unique Pathways** | **Non-Hallmark Unique Pathways** |
| --- | --- | --- | --- |
| let-7(b/c/e)-5p | Adipogenesis, Apical Junction, DNA Repair, E2F Targets, G2M Checkpoint, Heme Metabolism, IL2-STAT5 Signaling, Mitotic Spindle, mTORC1 Signaling, MYC Targets v1, MYC Targets v2, TNFA Signaling via NFKB, Unfolded Protein Response, UV Response Up | TGFB Receptor, Translation, Nonsense Mediated Decay, Splicing, Cell Cycle, GTPase Cycle, TF Binding, MAPK Signaling, Histone Modifying Activity, Catalytic Activity on Nucleic Acids | Heterochromatin, TP53 Regulation, Metabolism of Nucleotides, snRNA Transcription |
| miR-29(a/b/c)-3p | EMT, Glycolysis, Hypoxia, mTORC1 Signaling, MYC Targets v1, Myogenesis, UV Response Down | Translation, Mitotic G2-G2/M Phases, Cell Cycle, Kinase Binding, Histone Modifying Activity, Immune Protein Signaling | PDGF, Collagen, Extracellular Matrix, Scavenger Receptors, NCAM1 |
| miR-365(a/b)-3p | - | Transcription Regulator Activity, Nuclear Protein Containing Complex, Protein DNA Complex | N/A |
| miR-34a-5p, miR-320a | Not Significant | Not Significant | Not Significant |
| miR-30(a/b/d)-5p | Adipogenesis, Apical Junction, Apoptosis, Complement, E2F Targets, G2M Checkpoint, Heme Metabolism, Hypoxia, IL2-STAT5 Signaling, mTORC1 Signaling, MYC Targets v1, Oxidative Phosphorylation, UV Response Up, Xenobiotic Metabolism | Translation, GTPase Cycle, Cell Cycle, Immune Protein Signaling, NOTCH Signaling, Splicing, Phosphatase Binding, Anchoring Junction, Transcription Coregulators | Golgi Associated Vesicle Biogenesis, Mitochondrial Translation, Selective Autophagy, Calcium Ion Binding |
| miR-93-5p, miR-20a-5p, miR-106b-5p | Adipogenesis, Allograft Rejection, Apical Junction, Apoptosis, Complement, EMT, Estrogen Response Early, Estrogen Response Late, Fatty Acid Metabolism, Heme Metabolism, Hypoxia, IL2-STAT5 Signaling, IL6-JAK-STAT3 Signaling, Interferon Alpha Response, Interferon Gamma Response, Mitotic Spindle, mTORC1 Signaling, Myogenesis, p53 Pathway, PI3K-AKT-mTOR Signaling, Protein Secretion, TGFB Signaling, TNFA Signaling via NFKB | Autophagy, ALK Signaling, Immune Protein Signaling, GTPase Cycle, MAPK Signaling, Phosphatase Binding, Cell Adhesion, Protein Kinase Activity | DDX58 IFIH1, Phagophore Assembly |
| miR-92(a/b)-3p, miR-25-3p | Adipogenesis, Complement, Mitotic Spindle, Oxidative Phosphorylation, p53 Pathway, TNFA Signaling via NFKB, Unfolded Protein Response | Transcription, Vesicle Mediated Transport, Semaphorin Interactions, Translation | Ruffle, Cortical Cytoskeleton, Metabolism of Vitamins And Cofactors |
| miR-221-3p, miR-222-3p, miR-542-3p | Not Significant | Not Significant | Not Significant |
| miR-24-3p, miR-574-3p, let-7i-3p | Not Significant | Not Significant | Not Significant |
| miR-155-5p, miR-142-3p, miR-205-5p | Not Significant | Not Significant | Not Significant |
| miR-15a-5p, miR-103a-3p, miR-107 | DNA Repair, E2F Targets, G2M Checkpoint, Heme Metabolism, Mitotic Spindle, MYC Targets v1 | WNT Signaling, Protein Kinase Activity, Cell Adhesion, TGFB, Transcription Factor Binding | Autophagosome, Regulated Necrosis,, Beta Catenin Independent WNT Signaling, Transcriptional Regulation by RUNX1 |
| miR-125b-5p, miR-199a-5p, miR-199a-3p | E2F Targets, Glycolysis, IL2-STAT5 Signaling, mTORC1 Signaling, MYC Targets v1, MYC Targets v2, Oxidative Phosphorylation, PI3K-AKT-mTOR Signaling | PTEN Regulation, ALK Signaling, TP53 Regulation, Cell Cycle, NOTCH Signaling, Translation, Histone Modifying Action, GTPase Cycle, RTKs, Immune Protein Signaling | Transcription Corepressor Binding, Promoter Specific Chromatin Binding, DNA Binding |
| miR-484, miR-193b-3p, miR-26b-5p | DNA Repair, E2F Targets, G2M Checkpoint, Hypoxia, Mitotic Spindle, mTORC1 Signaling, MYC Targets v1, Oxidative Phosphorylation, p53 Pathway, Unfolded Protein Response | Translation, Nonsense Mediated Decay, Cell Cycle, Immune Protein Signaling, Cell Adhesion, Transcription Coregulator Activity, RTKs, Single Stranded DNA Binding | Cytosolic Large Ribosomal Subunit, DNA Helicase Activity, Transcription Factor Activation, Histone Methyltransferase Complex, Dioxygenase Activity, Sm-like Protein Family Complex |
| miR-16-5p, miR-423-5p, miR-424-4p | Not Significant | Not Significant | Not Significant |
| miR-10a-5p, miR-10b-5p, miR-140-3p | E2F Targets, G2M Checkpoint, Heme Metabolism, mTORC1 Signaling, MYC Targets v1, Oxidative Phosphorylation, UV Response Up | Robo Receptors, Translation, Nonsense Mediated Decay, WNT Signaling, Splicing, DNA Replication, Cell Cycle, NOTCH Signaling, Cell Adhesion, GTPase Cycle, Autophagy | Molecular Carrier Activity, Electron Transport, Translation Regulator Activity |
| miR-21-5p, miR-181a-5p, miR-27a-3p | E2F Targets, Fatty Acid Metabolism, G2M Checkpoint, Hypoxia, Mitotic Spindle, mTORC1 Signaling, MYC Targets v1, p53 Pathway, Peroxisome, PI3K-AKT-mTOR Signaling, Protein Secretion, Unfolded Protein Response, UV Response Down, UV Response Up | Sequence Binding, Translation, Toll-Like Receptor Cascades, Nonsense Mediated Decay, Immune Protein Signaling, DNA Secondary Structure Binding, MAPK Signaling, TP53 Regulation, RTKs, VEGF, ESR Signaling, Transcription Activity, Early Endosome, PKMTs Methylate Histone Lysines, Histone Modifying Activity, Vesicle Mediated Transport, TGFB, Cell Cycle, Adipogenesis, Apoptosis | Nuclear Localization Sequence Binding, RET Signaling, Pyruvate Metabolism, Insulin Receptor, ERBB2, TBC Rabgaps, DNA Damage |
| miR-23b-3p, miR-375, miR-23a-3p | Not Significant | Not Significant | Not Significant |
| miR-200c-3p, miR-324-5p, miR-182-5p | Apical Junction | ALK Signaling, RTKs, Endosome, GTPase Cycle, Vesicle Mediated Transport, Transcription Factor Binding, Immune Protein Signaling | Sarcolemma, Plasma Membrane Region |
| miR-100-5p, miR-99a-5p | Not Significant | Not Significant | Not Significant |

**Table S5.** Identified functions and top three miRNAs for the communities in the Her2 subtype network.

| **Top 3 miRNAs** | **Hallmark Pathways** | **Non-Hallmark, Non-Unique Pathways** | **Non-Hallmark Unique Pathways** |
| --- | --- | --- | --- |
| miR-106b-5p, miR-20a-5p, miR-93-5p | Adipogenesis, Androgen Response, Apoptosis, EMT, Estrogen Response Late, Heme Metabolism, Hypoxia, IL6-JAK-STAT3 Signaling, Interferon Gamma Response, Mitotic Spindle, mTORC1 Signaling, Myogenesis, PI3K-AKT-mTOR Signaling, Protein Secretion, TGFB Signaling, UV Response Down | ALK Signaling, RHO GTPase Cycle, STKs and RTKs, Toll-like Receptor Cascade, Immune Signaling, Cell Adhesion, SMAD | Phagophore Assembly Site, DDX58 IFIH1, TBC Rabgaps, Pyruvate Metabolism, PTK6 Signaling, TNF Signaling |
| miR-27(a/b)-3p, miR-128-3p | E2F Targets, G2M Checkpoint, Mitotic Spindle, mTORC1 Signaling, MYC Targets v1, MYC Targets v2, p53 Pathway, TNFA Signaling via NFKB, UV Response Down | Cell Cycle, Translation, Histone Methyltransferase Activity, MAPK Signaling, Chromatin Binding, Transcription Factor Binding, Immune Protein Signaling, Kinase Activity, Condensed Chromosome, Acyltransferase Activity, Apoptosis | Histone Methyltransferase Activity, PKMTs Methylate Histone Lysines, SH3 Domain Binding |
| miR-26b-5p, miR-181(a/b)-5p | Allograft Rejection, Apoptosis, Complement, DNA Repair, EMT, Estrogen Response Early, Estrogen Response Late, Hypoxia, IL2-STAT5 Signaling, Interferon Gamma Response, Mitotic Spindle, mTORC1 Signaling, MYC Targets v1, Oxidative Phosphorylation, TNFA Signaling via NFKB, Unfolded Protein Response, UV Response Down, UV Response Up | Translation, Immune Protein Signaling, Nonsense Mediated Decay, Vesicle Mediated Transport, Cell Cycle, Phosphatase Complex, Transcription Activity, MAPK Signaling, Kinase Activity, TGFB, TP53 Regulation | Sumoylation of Transcription Cofactors, COPII Mediated Vesicle Transport, BRAF and RAF1 Signaling |
| miR-193b-3p, miR-320a-3p, miR-378a-5p | Not Significant | Not Significant | Not Significant |
| miR-29(a/b/c)-3p | Apical Junction, Apoptosis, E2F Targets, EMT, Mitotic Spindle, MYC Targets v1, Myogenesis, PI3K-AKT-mTOR Signaling, UV Response Down | SMAD, Cell Cycle, Histone Modifying Activity, Immune Protein Signaling, Kinase Activity, Vesicle Mediated Transport, GTPase Cycle, MAPK Signaling | Collagen, MET, Extracellular Matrix, PDGF |
| miR-21-5p, miR-21-3p, miR-409-3p | Not Significant | Not Significant | Not Significant |
| miR-30(a/b/c)-5p | E2F Targets, Fatty Acid Metabolism, G2M Checkpoint, Glycolysis, Hypoxia, IL2-STAT5 Signaling, mTORC1 Signaling, MYC Targets v1, TNFA Signaling via NFKB, Unfolded Protein Response, UV Response Up | Cell Cycle, GTPase Cycle, Immune Protein Signaling, ALK Signaling, RTKs, Apoptosis, Autophagy, MAPK Signaling, DNA Repair, Mitotic Spindle | Far Sin Stripak Complex, RNA Exonuclease Activity, PKR Mediated Signaling, Selective Autophagy, Pre-NOTCH Expression and Processing |
| miR-484, miR-331-3p, miR-455-3p | Adipogenesis, DNA Repair, E2F Targets, Fatty Acid Metabolism, G2M Checkpoint, Glycolysis, Hypoxia, Interferon Gamma Response, mTORC1 Signaling, MYC Targets v1, Oxidative Phosphorylation, p53 Pathway, TNFA Signaling via NFKB, Unfolded Protein Response, UV Response Up | Translation, Nonsense Mediated Decay, NOTCH Signaling, Immune Protein Signaling, DNA Replication, MAPK Signaling, Cell Cycle, Kinase Activity, Cell Adhesion, Actin Cytoskeleton | Ribosomal Subunit, Translation Elongation Factor, Degradation of Beta Catenin, Regulation of RAS, Nucleotide Excision Repair, Interleukin Signaling |
| miR-103a-3p, miR-107, miR-200c-3p | Adipogenesis, Apical Junction, Apoptosis, Coagulation, Complement, EMT, G2M Checkpoint, Heme Metabolism, Mitotic Spindle, mTORC1 Signaling, p53 Pathway, Unfolded Protein Response | ALK Signaling, VEGF Signaling, Toll-like Receptor Cascades, RTKs, GTPase Cycle, Cell Cycle, Vesicle Mediated Transport, Phospholipid Binding, WNT Signaling | RAC3 GTPase, VEGF Signaling, Semaphorin Interactions |
| miR-92(a/b)-3p, miR-25-3p | Fatty Acid Metabolism, G2M Checkpoint, Heme Metabolism, Mitotic Spindle, p53 Pathway | Acyltransferase, Cell Adhesion, Cell Substrate Junction, GTPase Cycle | Beta Catenin Binding, Specific Granule |
| miR-218-5p | Not Significant | Not Significant | Not Significant |
| let-7(a/b/d)-5p | Adipogenesis, Apoptosis, DNA Repair, E2F Targets, MYC Targets v1, Oxidative Phosphorylation | Translation, Kinase Activity, Electron Transport, Vesicle Mediated Transport, Chromatin Binding | Protein STK Activator Activity, Heterochromatin, MITF-M |
| miR-155-5p, miR-26a-5p, miR-378a-3p | Not Significant | Not Significant | Not Significant |
| miR-16-5p, miR-15(a/b)-5p | TNFA Signaling via NFKB | Translation, Cell Substrate Junction, Cell Cycle, Phospholipid Binding | tRNA Processing, Gene Silencing by RNA |
| miR-23(a/b)-3p, miR-365-3p | E2F Targets, G2M Checkpoint, Oxidative Phosphorylation, TNFA Signaling via NFKB | PTEN Regulation, NOTCH Signaling, WNT Signaling, B-cell Receptor, Translation, TP53 Regulation, Cell Cycle, Splicing, Apoptosis, ESR Mediated Signaling, MAPK Signaling, Kinase Activity, RTKs | Cyclin A, Transcription Regulator Activity, ATPase Complex |
| miR-19(a/b)-3p, miR-335-3p | Glycolysis, Hypoxia, mTORC1 SIgnaling, TNFA Signaling via NFKB, Xenobiotic Metabolism | Cell Substrate Junction, Endosome, Vesicle Mediated Transport, Acyltransferase Activity | - |
| miR-10a-5p, miR-339-5p, miR-10b-5p | Not Significant | Not Significant | Not Significant |
| miR-24-3p, miR-34a-5p, miR-221-3p | Not Significant | Not Significant | Not Significant |
| miR-148a-3p, miR-148b-3p, miR-152-3p | Heme Metabolism, Oxidative Phosphorylation, p53 Pathway | HMOX1, Respiratory Electron Transport, Transport Vesicle, GTPase Cycle, Splicing, TP53 Regulation, Immune Protein Signaling | Synaptic Vesicle Membrane, Transporter Activity |
| miR-192-5p | Not Significant | Not Significant | Not Significant |

**Table S6.** Identified functions and top three miRNAs for the communities in the LumA subtype network.

| **Top 3 miRNAs** | **Hallmark Pathways** | **Non-Hallmark, Non-Unique Pathways** | **Non-Hallmark Unique Pathways** |
| --- | --- | --- | --- |
| miR-484, miR-16-5p, miR-455-3p | Apical Junction, DNA Repair, E2F Targets, EMT, G2M Checkpoint, Glycolysis, Heme Metabolism, Hypoxia, Mitotic Spindle, mTORC1 Signaling, MYC Targets v1, Myogenesis, Oxidative Phosphorylation, p53 Pathway, TGFB Signaling, TNFA Signaling via NFKB | Translation, GTPase Cycle, Ribosome, Cell Substrate Junction, Cell Adhesion, Adipogenesis, Interferon Gamma, PTEN Regulation | Translation Elongation Factor, Small Ribosomal Subunit, Sumo Transferase, rRNA Binding, FGFR2 Alternative Splicing, SMAD, Nuclease Activity, ADP Binding, Single Stranded RNA Binding, PTEN Gene Transcription, PTK6, Methyltransferase, Sm-like protein family |
| miR-15a-5p, miR-103a-3p, miR-195-5p | Apical Junction, Complement, G2M Checkpoint, Glycolysis, Heme Metabolism, Mitotic Spindle, mTORC1 Signaling, MYC Targets v1 | Antigen Processing, Acyltransferase, TGFB, Vesicle Mediated Transport, Cell Cycle, Phosphatase Binding, GTPase Cycle, ALK Signaling | PI Metabolism, ISG15 Antiviral Mechanism, Synapses Transmission |
| miR-375, miR-365(a/b)-3p | N/A | Transcription Regulator, Second Messengers, Phosphatase Activity, Cell Substrate Junction, RTKs, Actin Cytoskeleton, Endosome, Kinase Binding, Immune Signaling | N/A |
| miR-218-5p, miR-34a-5p, miR-320a | Not Significant | Not Significant | Not Significant |
| miR-30(a/b/c)-5p | Apical Junction, Apoptosis, Coagulation, Complement, EMT, Glycolysis, Hypoxia, Interferon Gamma Response, KRAS Signaling Up, Mitotic Spindle, mTORC1 Signaling, MYC Targets v1, p53 Pathway, Protein Secretion, TNFA Signaling via NRKB, UV Response Down, UV Response Up, Xenobiotic Metabolism | ALK Signaling, MAPK Signaling, GTPase Cycle, WNT Signaling, ESR Mediated Signaling, Protein Dimerization, Cell Adhesion | Euchromatin, RTKs, Peptidase Activity, Double Stranded RNA Binding, Apoptotic Execution Phase, Semaphorin Interactions, L1CAM Interactions, Ephrin Signaling, RUNX3, RUNX1 |
| miR-130a-3p, miR-148a-3p, miR-30a-3p | N/A | Splicing, Endosome, Post-translational Protein Modification, Microtubule, Vesicle Mediated Transport, GTPase Cycle, Nervous System Development | DNA Binding Transcription Factor Activity, DNA Binding Transcription Repressor Activity |
| miR-23-3p, miR-142-3p | Not Significant | Not Significant | Not Significant |
| miR-29(a/b/c)-3p | Apical Junction, Apoptosis, EMT, MYC Targets v1, Myogenesis, UV Response Down | Ubiquitin Ligase Complex, MAPK Signaling, Acyltransferase, Immune Protein Signaling | PDGF, Collagen, Extracellular Matrix, MET, Scavenger Receptors, NCAM1, Calcium Ion Binding |
| let-7(a/b/c)-5p | Adipogenesis, Androgen Response, Apical Junction, Apoptosis, DNA Repair, E2F Targets, G2M Checkpoint, IL2-STAT5 Signaling, MYC Targets v1, MYC Targets v2, Oxidative Phosphorylation, TNFA Signaling via NFKB, Unfolded Protein Response, UV Response Up | Translation, Nucleotide Excision Repair, rRNA, Splicing, Cell Cycle, WNT Signaling | rRNA Modification, MITF-M, Single Stranded DNA Binding, DNA Double Strand Break Repair |
| miR-155-5p, miR-150-5p, miR-24-3p | Not Significant | Not Significant | Not Significant |
| miR-92a-3p, miR-21-5p, miR-25-3p | Not Significant | Not Significant | Not Significant |
| miR-125b-5p, miR-10a-5p, miR-22-3p | Adipogenesis, DNA Repair, mTORC1 Signaling, MYC Targets v1, Oxidative Phosphorylation | Splicing, Nonsense Mediated Decay, ESR Signaling, Translation, PTEN Regulation, Estrogen Dependent Gene Expression, Cell Cycle, Ribonucleoprotein Complex Binding, Cell Adhesion, Phosphatase Activity, Apoptosis, MAPK Signaling, TP53 Regulation | Transcription Corepressor Binding, mRNA Splicing, Protein Serine Threonine Phosphatase Activity, , Mitotic G1 Phase and G1/S Transition, Cyclin A |
| miR-181(a/b)-5p, miR-27a-3p | Apoptosis, Estrogen Response Early, Estrogen Response Late, G2M Checkpoint, Mitotic Spindle, mTORC1 Signaling, MYC Targets v1, p53 Pathway, PI3K-AKT-mTOR Signaling, Unfolded Protein Response, UV Response Down | Histone Activity, Cell Cycle, PTEN Regulation, p53 Binding, Transcription Factor Binding, RTKs, TGFB, MITF-M | PKMTs, GDP Binding, Kinase Activator Activity |
| miR-26a-5p, miR-221-3p, miR-100-5p | Adipogenesis, E2F Targets, G2M Checkpoint, Glycolysis, Hypoxia, mTORC1 Signaling, MYC Targets v1, Oxidative Phosphorylation | Nucleosome, Peptidase Complex, Cell Cycle, PTEN Regulation, MAPK Signaling, GTPase Cycle, B-Cell Receptor, Immune Protein Signaling, SMAD, Enzyme Activity, Autophagosome, Molecular Adaptor Activity, Phosphatase Binding, DNA Damage, Apoptosis, NOTCH Signaling | Nucleosomal Binding, Ubiquitination, ABC Family, DNA Replication, Degradation of Beta Catenin, Electron Transfer |
| miR-93-5p, miR-20a-5p, miR-106b-5p | Adipogenesis, Apoptosis, Interferon Gamma Response, Mitotic Spindle, MYC Targets v1, p53 Pathway, Protein Secretion, TGFB Signaling, TNFA Signaling via NFKB, UV Response Down | MAPK Signaling, ALK Signaling, Immune Protein Signaling, GTPase, NOTCH Signaling, Cell Adhesion, TP53 Regulation | Toll-like Receptor Cascade, Interleukin Signaling, Senescence, Mitochondrial Biogenesis, FOXO, DDX58 IFIH1 |
| miR-361-5p | Not Significant | Not Significant | Not Significant |

**Table S7.** Identified functions and top three miRNAs for the communities in the LumB subtype network.

| **Top 3 miRNAs** | **Hallmark Pathways** | **Non-Hallmark, Non-Unique Pathways** | **Non-Hallmark Unique Pathways** |
| --- | --- | --- | --- |
| let-7(a/b)-5p, miR-98-5p | Adipogenesis, Androgen Response, DNA Repair, E2F Targets, MYC Targets v1, Oxidative Phosphorylation, p53 Pathway, Protein Secretion, UV Response Up | rRNA, Translation, Nucleotide Excision Repair, Vesicle Mediated Transport, Nonsense Mediated Decay, Splicing, Immune Protein Signaling, WNT Signaling, PTEN Regulation, Cell Cycle, DNA Damage, MITF-M | Mitochondrial Biogenesis, SWI/SNF Superfamily, Heterochromatin, Ubiquitin Binding, FGFR, Electron Transfer, Telomere Maintenance |
| miR-186-5p, miR-181(a/b)-5p | Androgen Response, Estrogen Response Early, mTORC1 Signaling, MYC Targets v1, Oxidative Phosphorylation, UV Response Up | Nonsense Mediated Decay, Translation, PTEN Regulation, rRNA, Estrogen Dependent Gene Expression, Splicing, Cell Cycle, Toll-like Receptor Cascade, WNT Signaling, MAPK Signaling, Autophagy, Enzyme Activity, Chromatin Binding, Apoptosis, Oxidoreductase, DNA Repair, Interferon Signaling, RKTs | Nuclear Envelope Reassembly, Small Ribosomal Subunit, UCH Proteinases, Kinase Activator Activity, Nucleosome Binding, Spindle Microtubule, Senescence, DNA Replication Pre-initiation |
| miR-484, miR-331-3p, miR-125-5p | Adipogenesis, Allograft Rejection, Apoptosis, Complement, Glycolysis, Hypoxia, IL2-STAT5 Signaling, Mitotic Spindle, mTORC1 Signaling, MYC Targets v1, MYC Targets v2, Myogenesis, Oxidative Phosphorylation, PI3K-AKT-mTOR Signaling, Unfolded Protein Response | Transcription, Translation, RHO GTPase Cycle, mRNA Splicing, Protein Serine Threonine Kinase Activity, Immune Protein Signaling, Cell Adhesion, Cell Substrate Junction, Microtubule, RKTs | Transcription Corepressor Binding, Citric Acid Cycle, Cytosolic Sensors of Pathogen Associated DNA, Spliceosome, p75 NTR |
| miR-193b-3p, miR-221-3p, miR-222-3p | Not Significant | Not Significant | Not Significant |
| miR-24-3p, miR-148a-3p, miR-196a-5p | Not Significant | Not Significant | Not Significant |
| miR-26b-5p, miR-27(a/b)-3p | Adipogenesis, Apoptosis, Mitotic Spindle, mTORC1 Signaling | Vesicle Mediated Transport, TP53 Regulation, Histone Activity, DNA Damage, DNA Repair, Cell Adhesion, Acyltransferase, Cell Cycle, Unfolded Protein Response | PKMTs, Methyltransferase Activity, Damaged DNA Binding, Protein Ubiquitination, Cell-Cell Communication |
| miR-155-5p, miR-21-5p, miR-140-3p | Not Significant | Not Significant | Not Significant |
| miR-375, miiR-342-3p, miR-200a-3p | E2F Targets, EMT, Hypoxia, mTORC1 Signaling, MYC Targets v1, Oxidative Phosphorylation, PI3K-AKT-mTOR Signaling, Unfolded Protein Response | NRTKs, ALK Signaling, Cell Substrate Junction, Apoptosis, Transcription Factor Binding, Chromatin BInding, Senescence, RHO GTPase Cycle, MAPK Signaling, Immune Protein Signaling | FCGR, Protein Tyrosine Kinase Binding, SH3 Domain Binding, Phosphoric Ester Hydrolase Activity, Calcium Ion Binding |
| miR-106b-5p, miR-20a-5p, miR-93-5p | Adipogenesis, Apical Junction, Apoptosis, Complement, Fatty Acid Metabolism, Heme Metabolism, Interferon Alpha Response, Interferon Gamma Response, Oxidative Phosphorylation, Protein Secretion, Unfolded Protein Response | ALK Signaling, GTPase Cycle, Vesicle Mediated Transport, Toll-Like Receptor Cascade, Cell Cycle, Cell Adhesion, Acyltransferase, Cell Substrate Junction, Transcription Factor Binding, TGFB, MITF-M, TP53 Regulation, WNT Signaling, MAPK Signaling, DNA Damage | MECP2, DDX58 IFIH1, Actomyosin, MET, Necrosis, Interleukin Signaling |
| miR-15b-3p | Not Significant | Not Significant | Not Significant |
| miR-15(a/b)-5p, miR-195-5p | Apical Junction, Apoptosis, Coagulation, Complement, EMT, G2M Checkpoint, Glycolysis, Heme Metabolism, Hypoxia, Mitotic Spindle, mTORC1 Signaling, MYC Targets v1, Oxidative Phosphorylation, p53 Pathway, PI3K-AKT-mTOR Signaling, Protein Secretion, TGFB Signaling, TNFA Signaling via NFKB, Unfolded Protein Response, UV Response Down | GTPase Cycle, Toll-like Receptor Cascade, VEGF, Immune Protein Signaling, B-Cell Receptor, Cell Cycle, Phosphatase Binding, Cell Substrate Junction, Cell Adhesion, Phospholipid Binding, MAPK Signaling, WNT Signaling, NOTCH Signaling | Autophagosome, SMAD, G Protein Activity, PI Metabolism, Sarcoplasm, Platelet Alpha Granule, Protein Serine Threonine Phosphatase, TCR Signaling, |
| miR-34a-5p, miR-320a, miR-10-5p | Not Significant | Not Significant | Not Significant |
| miR-92a-3p, miR-25-3p, miR-500a-3p | Not Significant | Not Significant | Not Significant |
| miR-30(a/b/c)-5p | Apical Junction, Complement, DNA Repair, Fatty Acid Metabolism, G2M Checkpoint, Glycolysis, Heme Metabolism, mTORC1 Signaling, MYC Targets v1, Oxidative Phosphorylation, Protein Secretion, Unfolded Protein Response, UV Response Up, Xenobiotic Metabolism | rRNA, ALK Signaling, DDX58 IFIH1, Nucleotide Excision Repair, Vesicle Mediated Transport, WNT Signaling, GTPase Cycle, Cell Cycle, MAPK Signaling, NOTCH Signaling, Immune Protein Signaling, Apoptosis, DNA Replication, B-Cell Receptor, Phospholipid Binding, Cell Adhesion, ESR Mediated Signaling, Protein Dimerization, Autophagy, RTKs | Phosphatase Complex, Phosphatidylinositol Bisphosphate Binding, S Phase, DNA Synthesis, Mitotic Prometaphase |
| miR-183-5p | Not Significant | Not Significant | Not Significant |
| miR-455-3p, miR-92b-3p, miR-149-5p | DNA Repair, Interferon Gamma Response, Mitotic Spindle, mTORC1 Signaling, MYC Targets v1, MYC Targets v2, Oxidative Phosphorylation, p53 Pathway | Translation, NOTCH Signaling, WNT Signaling, PTEN Regulation, Splicing, Immune Protein Signaling, Endosome, Hydrolysis, Transcription Factor Binding, Cell Adhesion, RHO GTPase Cycle, Cell Substrate Junction, Protein Dimerization, MAPK Signaling, RKTs | Peoptidase Complex, Organic Acid Binding, Carbohydrate Binding |
| miR-29(a/b/c)-3p | E2F Targets, EMT, G2M Checkpoint, MYC Targets v1, Myogenesis | Vesicle Mediated Transport, DNA Repair, Dioxygenase Activity, Apoptosis, Histone Modifying Activity, Cell Cycle, Immune Protein Signaling |  |
